# Axolotl regeneration reveals a dormant cis-regulatory grammar conserved across vertebrate genomes

**DOI:** 10.64898/2026.07.13.738357

**Authors:** Takashi Fujiwara, Kazuki Nakanishi, Takahide Suzuki, Hideyuki Shimizu

## Abstract

Unlike most mammals, which lack the capacity to regenerate complex tissues following injury, other vertebrates such as the axolotl rebuild complete limbs throughout life, yet the regulatory mechanisms underlying this striking difference have remained elusive. Here, we define the core cis-regulatory motif grammar driving axolotl limb regeneration and demonstrate that this grammar is conserved within the syntenic neighborhoods of regeneration-gene orthologs in human and mouse genomes, despite being epigenetically sealed in adult mammalian tissues. Integrating this cross-species grammar projection with AlphaGenome, a multimodal genomic AI capable of predicting epigenomic states from long sequence context, we find that the highest-ranking candidate loci are predicted to occupy a state of bivalent dormancy marked by the co-enrichment of poising and repressive histone modifications alongside suppression of transcriptional activity and chromatin accessibility. Systematic *in silico* motif perturbation further predicts that this dormant state is actively enforced by specific dormancy-stabilizing sequence elements, and that disrupting these elements shifts candidate loci toward a regeneration-competent chromatin configuration. Our findings support a model in which the regenerative blueprint has not been erased from the mammalian genome but locked within it, opening new avenues for understanding the evolution of regenerative competence and for the rational reactivation of latent regenerative programs.

## Introduction

The loss of regenerative capacity in adult mammals remains one of the most profound mysteries in regenerative medicine and evolutionary developmental biology^1–4^. Regeneration-competent vertebrates such as axolotls and zebrafish restore amputated appendages, injured hearts, and damaged nervous tissue with remarkable fidelity^4–7^, whereas comparable injuries in adult mammals typically culminate in fibrosis, scarring, and incomplete functional recovery^1,4^. This contrast has long been interpreted as an evolutionary loss of regenerative potential, yet the molecular components required for tissue growth, patterning, and repair remain broadly represented in mammalian genomes. Most strikingly, the neonatal mouse’s heart regenerates fully after injury at postnatal day 1 but loses this capacity irreversibly within the first postnatal week as tissue maturation proceeds^8,9^. That mammals retain a functional regenerative window in early life argues that the deficit in adult regeneration reflects not the absence of regenerative genes but the failure to deploy an otherwise latent regulatory program.

Regenerative responses are orchestrated in large part through tissue regeneration enhancer elements (TREEs), cis-regulatory sequences that couple tissue injury to the coordinated activation of developmental and repair-associated transcriptional programs^10–13^. TREE activity is defined not by a single universal sequence but by combinatorial transcription-factor motif grammar, meaning the arrangement and combination of short DNA motifs recognized by transcription factors that integrate injury-responsive signaling, developmental patterning, and cellular state. Among regeneration-competent vertebrates, the axolotl offers a particularly informative system for defining this cis-regulatory logic: it retains the capacity to regenerate complete limbs throughout adult life, and limb amputation triggers a stereotyped sequence of wound closure, blastema formation, outgrowth, and redifferentiation. During this process, injury-responsive signaling is coordinated with developmental patterning and progenitor-like cellular states, making the regenerating limb a useful setting in which to infer the regulatory grammar of productive vertebrate regeneration^14–17^. Previous studies have identified injury-responsive regeneration enhancers in regeneration-competent animals^10–13^ and have shown that some injury-responsive regulatory programs become silenced as animals mature^8,9^. These findings suggest that loss of regeneration can occur through regulatory silencing, but they do not resolve whether mammals have lost regeneration-associated cis-regulatory grammar itself or instead retain such grammar near orthologous gene neighborhoods in a non-deployed, dormant chromatin state. Here, a dormant chromatin state refers to a predicted enhancer-like sequence context that lacks active-enhancer deployment and chromatin accessibility.

Identifying mammalian loci that retain regenerative grammar presents two compounding challenges. First, enhancer sequences undergo rapid evolutionary turnover, meaning that direct alignment of axolotl regeneration-associated elements to mammalian genomes recovers only a limited subset of functionally analogous loci^18–20^. Second, even when regeneration-associated motifs are present in mammalian genomes, their mere existence does not establish that the locus is epigenetically competent to respond to injury, as the same motif grammar can reside within active chromatin, poised chromatin, polycomb-repressed chromatin associated with H3K27me3, or inaccessible chromatin. Resolving the chromatin state of motif-containing loci requires integrating sequence information across genomic scales that exceed the reach of conventional motif-scanning and short-window sequence analysis. No framework has yet addressed these requirements jointly, namely cross-species grammar projection, chromatin-state prediction, and functional sensitivity to sequence perturbation at candidate regenerative loci.

Here, we develop an axolotl-to-mammal framework that addresses these challenges by transferring regenerative regulatory information at the level of orthologous gene neighborhoods and transcription-factor motif grammar, bypassing the barrier of primary sequence divergence (**Fig. 1**). Integrating stage-resolved chromatin-accessibility and single-cell transcriptomic profiles from axolotl limb regeneration with AlphaGenome^21^, a multimodal genomic AI capable of integrating long sequence context into epigenomic predictions, we chart the regulatory landscape of 649 candidate loci across the human and mouse genomes and show that the highest-scoring regions are predicted to occupy a previously uncharacterized state of bivalent dormancy, supported by comparisons with matched genomic backgrounds and human disease-variant annotations. We refer to these computationally nominated mammalian intervals as dormant regeneration enhancer candidates (DRECs): candidate regulatory loci near regeneration-gene orthologs that retain axolotl-derived regenerative motif grammar but are predicted to lack active-enhancer deployment. Systematic perturbation of nearly 80,000 motif instances across human-mouse matched orthologous loci indicates that this dormant state is actively enforced by specific dormancy-stabilizing sequence elements whose disruption is predicted to shift candidate loci toward a regeneration-competent chromatin configuration. The recovery of these repressive elements in both human and mouse orthologous neighborhoods, despite an estimated 350 million years of divergence^22^, supports a model in which the regenerative deficit of adult mammals reflects regulatory sequestration rather than genetic erasure of regenerative potential.

**Figure 1.**
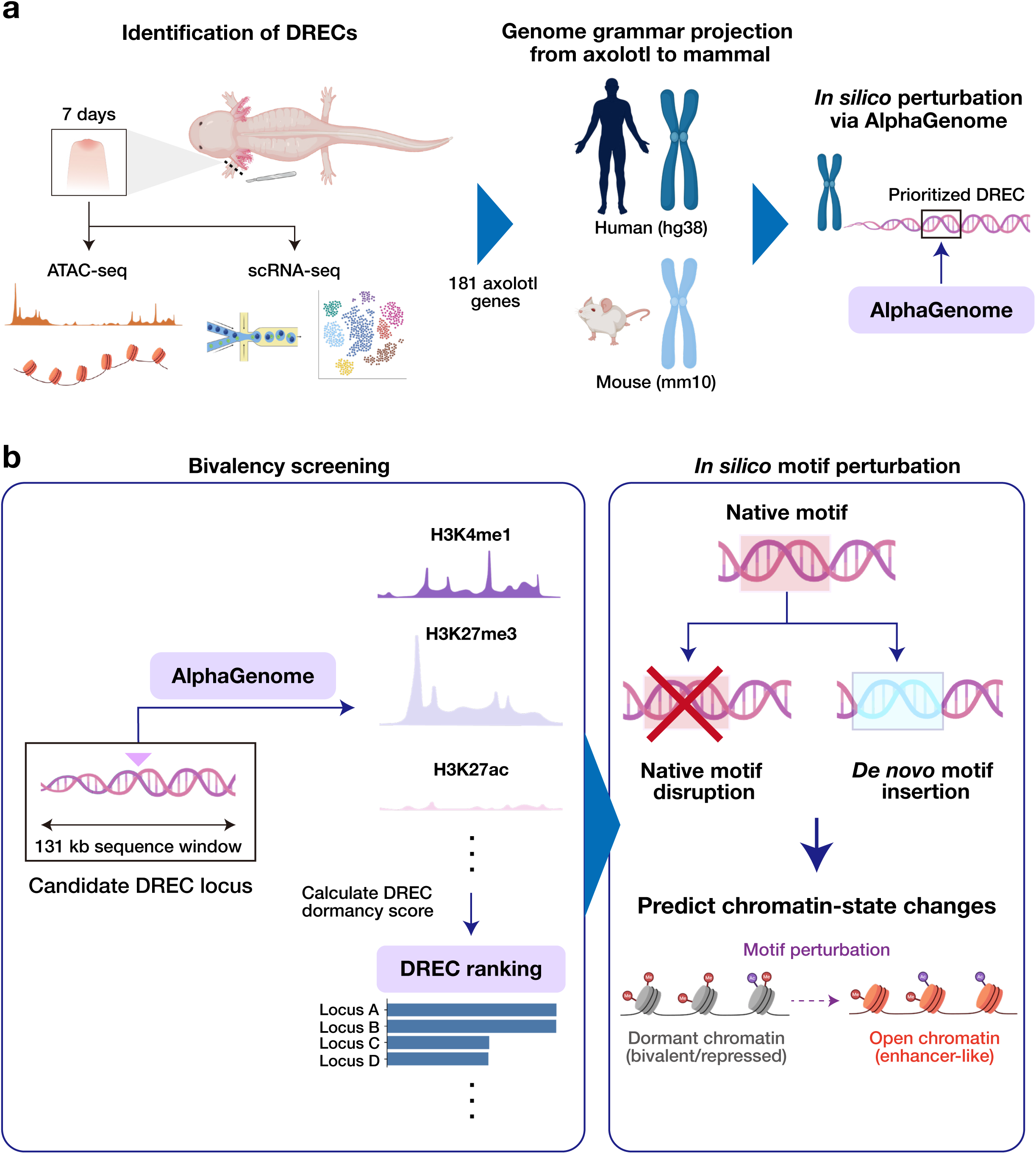
An axolotl-to-mammal computational framework for identifying dormant regeneration enhancer candidates (DRECs). **(a)** Overview of the cross-species DREC discovery framework. Stage-resolved ATAC-seq and single-cell RNA-seq (scRNA-seq) profiles across 0 h to 33 d of axolotl limb regeneration were integrated at the peak-gene level, and the day-7 mid-bud state was used to define a focused 181-gene regenerative prior. Regeneration-associated genes and their corresponding cis-regulatory motif grammar were projected onto orthologous gene neighborhoods in the human (hg38) and mouse (mm10) genomes. Candidate mammalian loci were evaluated by AlphaGenome-based *in silico* epigenomic scoring and motif-level perturbation to nominate putative DRECs. **(b)** AlphaGenome-based prioritization and functional interrogation of candidate loci. Left, bivalency screening: each candidate locus was evaluated within a 131 kb genomic sequence window, and predicted chromatin features, including H3K4me1, H3K27me3, and H3K27ac, were integrated into a composite DREC dormancy score used to rank candidate loci. Right, *in silico* motif perturbation: prioritized loci were subjected to native motif disruption (replacement with randomized sequences in triplicate) or *de novo* motif insertion, and the resulting changes in predicted chromatin accessibility and enhancer-associated histone modification profiles were quantified using AlphaGenome. Loci exhibiting a predicted transition from a dormant, bivalent chromatin state toward a more accessible, enhancer-like configuration were prioritized for further analysis.

## Results

### Stage-resolved multi-omic profiling defines the cis-regulatory grammar of axolotl limb regeneration

To construct a biologically grounded regulatory prior for mammalian DREC discovery, we reprocessed and integrated published stage-resolved chromatin-accessibility and single-cell gene-expression datasets spanning eight stages of axolotl limb regeneration: 0 h, 3 h, and days 1, 3, 7, 14, 22, and 33 (**Fig. 2a**). The number of accessible regions increased from 18,359 at 0 h to 36,371 at day 7, the largest number observed at any individual time point, and pooled peak calling across post-injury stages identified 67,489 non-redundant accessible regions (**Fig. 2b**). Of these, 15,132 were classified as regeneration-associated post-injury accessible regions, defined as peaks absent at 0 h but detected in at least two post-injury stages. Clustering of these regions identified transient, delayed, and sustained accessibility programs, including a prominent module peaking around day 7, whereas principal component analysis separated later regenerative stages from pre-injury and early-response states (**Supplementary Fig. 1a, b**). These results indicate that limb regeneration proceeds through a temporally organized succession of chromatin states.

**Figure 2.**
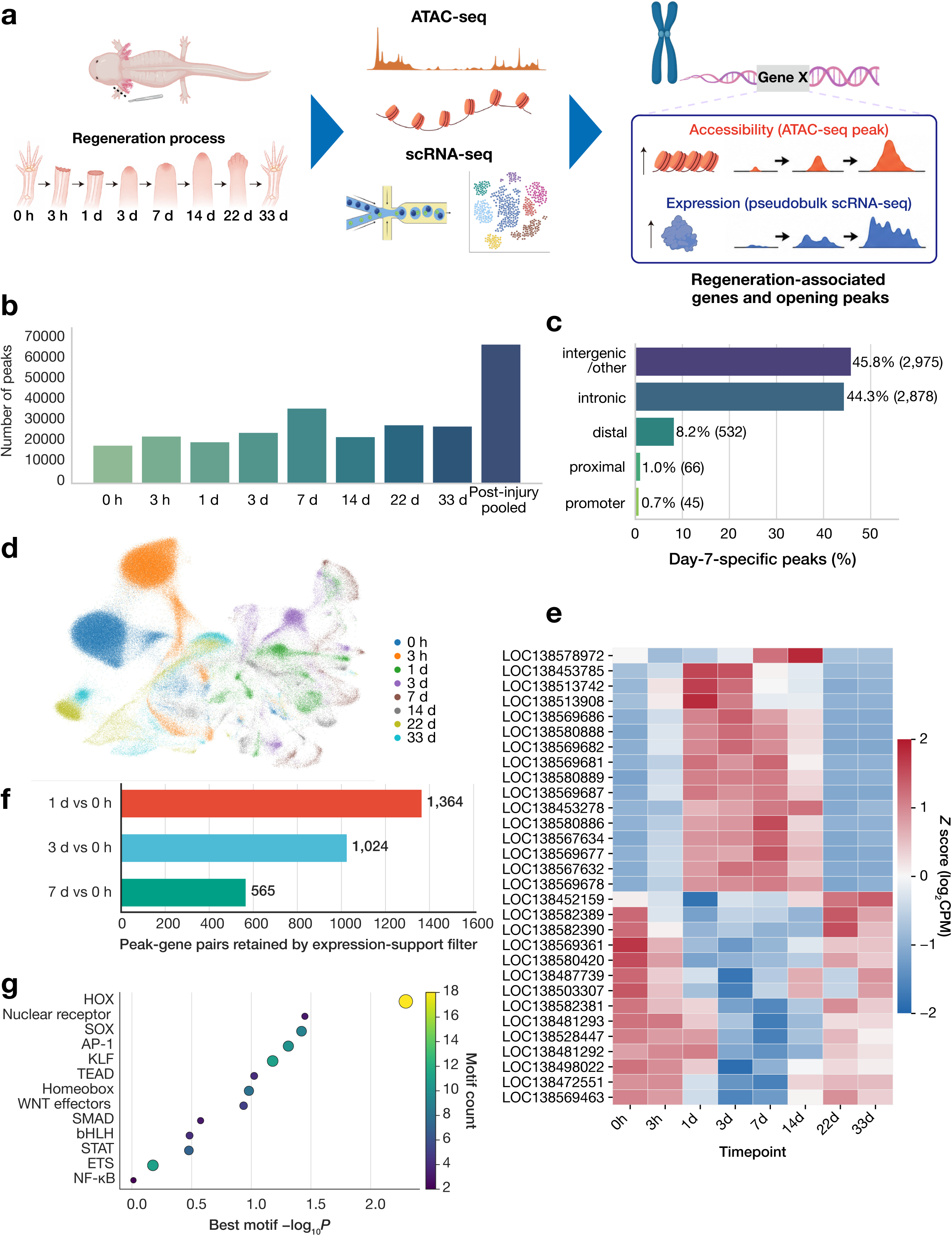
Stage-resolved multi-omic profiling defines the regenerative regulatory grammar of axolotl limb regeneration. **(a)** Experimental and analytical overview. Chromatin accessibility and transcriptional states were profiled by ATAC-seq and single-cell RNA-seq (scRNA-seq), respectively, across eight stages of axolotl limb regeneration: 0 h, 3 h, and days 1, 3, 7, 14, 22, and 33. Stage-matched accessibility and pseudobulk expression changes were integrated at the peak-gene level to identify regeneration-associated genes and dynamically opening regulatory elements. **(b)** Numbers of accessible chromatin regions detected at each stage. Peaks pooled across the seven post-injury stages comprised 67,489 non-redundant regions. **(c)** Genomic annotation of ATAC-seq peaks specifically accessible at the mid-bud stage (day 7). Bars show mutually exclusive peak_annotation_class assignments used for downstream support scoring: intergenic/other, non-exonic intronic gene-body, distal, proximal, and promoter-associated. These operational classes are distinct from raw feature-overlap annotations; exon-overlap calls were not plotted as an independent category in this panel. The predominance of intergenic/other (45.8%) and intronic (44.3%) peaks supports broad remodeling of putative distal enhancer landscapes during blastema formation. **(d)** UMAP visualization of single-cell transcriptomes collected across the eight-stage regeneration time course. Cells are colored according to sampling time point, illustrating the pronounced stage-dependent redistribution of transcriptional states. **(e)** Temporal expression dynamics of representative FDR-ranked expression-support genes (top 30 by adjusted *P* value; 16 induced and 14 repressed) during limb regeneration. Rows represent individual genes, ordered by the time point of maximal *Z* score, and columns represent time points (0 h through 33 d). Gene identifiers beginning with LOC denote unannotated axolotl gene models. Color scale indicates row-wise *Z* scores of log_2_-transformed pseudobulk counts per million (log_2_CPM). **(f)** Numbers of proximity-assigned ATAC-RNA peak-gene pairs retained through the FDR-thresholded expression-support filter at days 1, 3, and 7 relative to the uninjured 0-h state (1,364, 1,024, and 565 peak-gene pairs, respectively). The day-7 set was used as a focused mid-bud blastema prior for downstream cross-species projection. **(g)** Transcription factor (TF) motif families represented in the regeneration motif prior. The horizontal axis indicates the strongest motif-enrichment significance within each family, expressed as −log_10_ *P*, and point color and size indicate the number of enriched motifs assigned to the corresponding family. Represented families span developmental-patterning regulators (HOX, SOX, homeobox, WNT effectors), injury-and signal-responsive regulators (AP-1, STAT, NF-κB, SMAD), and additional factors (nuclear receptor, KLF, TEAD, bHLH, ETS), collectively defining a combinatorial regenerative grammar.

Because early time points preferentially capture wound-healing and inflammatory responses rather than regeneration-specific programs^7,23^, we prioritized day 7 for downstream candidate definition while retaining earlier stages for temporal comparison. At day 7, the blastema is established^14,24^, and the number of accessible regions reaches its maximum, making this stage a selective representation of the mid-bud regenerative state. To determine whether accessibility gained at this stage reflected promoter-proximal activation or broader regulatory remodeling, we assigned day-7-specific peaks to a mutually exclusive integration-region class used for downstream support scoring. Under this operational scheme, peaks were assigned primarily to intergenic/other (45.8%; 2,975 peaks) or non-exonic intronic gene-body classes (44.3%; 2,878 peaks), with a further 8.2% classified as distal (532 peaks), 1.0% as proximal (66 peaks), and 0.7% as promoter-associated (45 peaks) (**Fig. 2c**). This distribution supports broad remodeling of non-promoter regulatory landscapes during blastema formation.

In parallel, scRNA-seq revealed a stage-dependent redistribution of cellular transcriptional states (**Fig. 2d**). To enable stage-matched integration with bulk ATAC-seq while reducing cell-level sampling noise, we aggregated scRNA-seq counts by time point and performed pseudobulk differential-expression analysis. This analysis resolved early, intermediate, and late gene-expression programs (**Fig. 2e**), indicating that regeneration involves transcriptionally distinct phases rather than a uniform injury response. Integration of stage-matched ATAC-seq and single-cell-derived pseudobulk expression profiles retained 1,364, 1,024, and 565 proximity-assigned peak-gene pairs associated with genes passing the FDR-thresholded expression-support filter at days 1, 3, and 7 relative to 0 h, respectively (**Fig. 2f**). Because the single-cell transcriptomic dataset contained one library per time point, these links were used as a filtering and ranking prior for cross-species DREC discovery. The smaller day-7 set provided a focused mid-bud blastema prior for downstream cross-species projection. The day-7 associations involved 181 unique genes, which served as gene-level anchors for mammalian DREC discovery (**Fig. 1a**; **Supplementary Table 1**).

We next assessed the robustness and temporal coherence of the day-7 peak-gene associations. Linked genes included both induced and repressed transcripts, indicating that the integration was not restricted to strongly activated genes, and support-score ranking identified a set of high-confidence peak-gene pairs (**Supplementary Fig. 1c, d**). Stage-resolved ATAC-seq and RNA-seq profiles of the top-linked genes showed gene-specific temporal patterns, including both concordant and temporally offset changes in accessibility and expression (**Supplementary Fig. 2a**). These results support the peak-gene map as a temporally structured representation of the regenerative regulatory program.

Because individual enhancer sequences are poorly conserved across deep vertebrate divergence^18–20^ and the axolotl-mammal split spans approximately 350 million years^22^, orthologous gene identity provides a genomic anchor but is insufficient to transfer cis-regulatory logic across this evolutionary distance. We therefore represented the sequence-level features of regeneration-associated accessible regions as a transcription-factor motif grammar. Motif-enrichment analysis identified a broad combinatorial program involving developmental-patterning regulators, including HOX^25^, SOX^26^, homeobox^27^, and WNT-effector families^28,29^; injury- and signal-responsive regulators, including AP-1^30^, STAT^31^, NF-κB^32^, and SMAD families^33^; and additional nuclear-receptor^34^, KLF^35^, TEAD^36^, bHLH^37^, and ETS factors^38^ (**Fig. 2g**). This combinatorial architecture is consistent with a regulatory grammar that integrates developmental positional information with injury-responsive signaling.

The same day-7 expression-linked analysis, examined at the level of individual motifs rather than curated families, identified enrichment of AP-2γ^39^, HOXA13^40^, HIC1^41^, HOXC13^42^, and HOXA9^25^ motifs, together with additional developmental and signal-responsive factors (**Supplementary Fig. 2b, c**). Independent *de novo* motif discovery further recovered recurrent sequence patterns without relying on existing motif annotations (**Supplementary Fig. 2d**). Eligibility for insertion was defined before AlphaGenome outcome evaluation. Within the eligible primary-motif subset, motifs were ranked by decreasing HOMER-derived source_log_p_support, and the top three were selected: denovo_035 (HOMER header rank 24; CATGCAGATAAA; closest known match, FUS3), denovo_026 (header rank 15; TAGAATTGTGAA; closest known match, ZNF35), and denovo_033 (header rank 22; GCACTGAGAAAG; closest known match, PRDM1). These three motifs are distinct from the five *de novo* motifs displayed in **Supplementary Fig. 2d**. Together, the 181 regeneration-associated genes and their associated motif repertoire constitute an axolotl-derived regulatory prior for nominating motif-compatible mammalian candidate intervals for subsequent chromatin-state prediction.

### Cross-species projection and AlphaGenome scoring reveal that regenerative grammar is preserved but epigenetically sequestered in mammalian genomes

To investigate whether the axolotl regenerative grammar is preserved within human and mouse orthologous gene neighborhoods, we projected the 181 regeneration-associated genes onto their mammalian orthologs and scanned ortholog-centered genomic windows for compatible motif grammar, retaining clustered motif-positive intervals with multi-motif support for AlphaGenome-based chromatin-state scoring (**Fig. 3a**; **Materials and Methods**). This strategy transfers regulatory information at the level of conserved gene context and TF motif grammar rather than primary sequence alignment, thereby circumventing the limited enhancer sequence conservation across the approximately 350-million-year divergence separating axolotl and mammals^22^.

**Figure 3.**
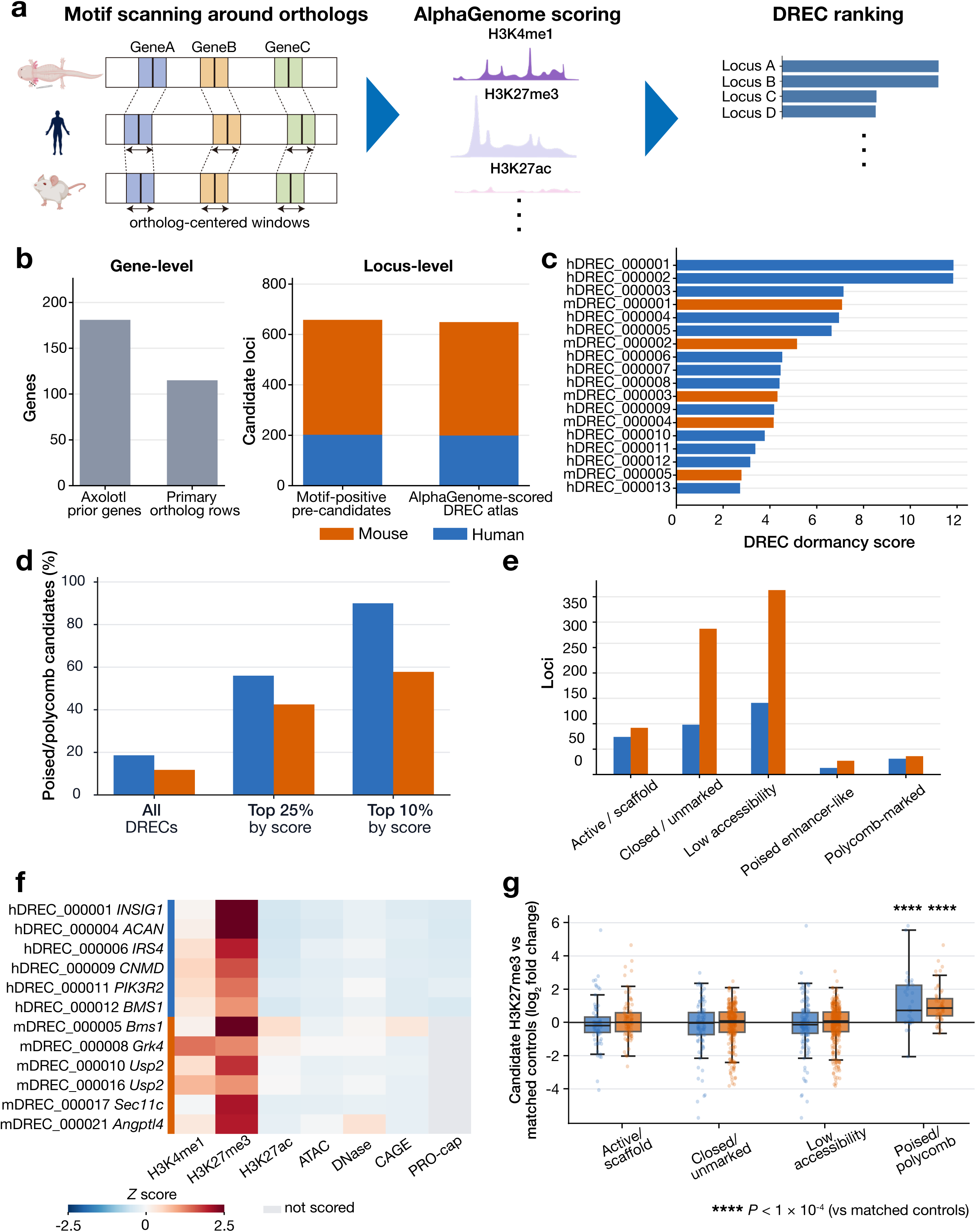
Cross-species projection, AlphaGenome scoring and candidate-level external support define a dual-species DREC atlas. **(a)** Schematic of the cross-species DREC nomination pipeline. Axolotl regeneration-associated genes were mapped to human and mouse orthologs, and orthologous genomic windows were scanned for regeneration-associated motif grammar. Motif-positive pre-candidates were evaluated using AlphaGenome to generate a composite DREC dormancy score, integrating predicted H3K4me1, H3K27me3, H3K27ac, and additional chromatin features. Candidate loci were ranked by this score. **(b)** Gene-level and locus-level summary of the DREC atlas. Left (Gene-level), numbers of axolotl prior genes (181) and primary ortholog rows used for cross-species projection (115). Right (Locus-level), stacked bars comparing motif-positive pre-candidate loci (202 human and 456 mouse; 658 total) with the AlphaGenome-scored DREC atlas (199 human and 450 mouse; 649 total), for human (blue) and mouse (orange) genomes. Nine low-information short motif clusters were excluded owing to insufficient unique motif support. **(c)** DREC dormancy scores for the top-ranked human (blue) and mouse (orange) DREC candidates. Candidates are ranked by score, with the highest-scoring loci shown at the top. Human loci are prefixed hDREC and mouse loci mDREC. **(d)** Enrichment of poised/polycomb annotations among high-scoring DRECs. Bars show the percentage of candidates assigned to poised-enhancer-like or polycomb-marked chromatin classes among all DRECs, the top 25% of candidates by DREC dormancy score, and the top 10% of candidates by score, separately for human and mouse. The increasing fraction among top-ranked candidates indicates that the DREC dormancy score preferentially prioritizes poised/polycomb-like loci. **(e)** Distribution of candidate loci by predicted chromatin class across human and mouse genomes. Chromatin classes include active/scaffold, closed/unmarked, low-accessibility, poised-enhancer-like, and polycomb-marked states. Class labels are non-exclusive; therefore, class-count totals can exceed the number of unique candidate loci. Thirteen candidates were labeled unclassified (3 human and 10 mouse); they are retained in **Supplementary Table 3** but are not plotted in this panel. **(f)** Heatmap of AlphaGenome-predicted epigenomic signals across representative DREC candidates selected from the poised-enhancer-like and/or polycomb-marked subset. Rows represent selected high-priority human and mouse candidates with linked gene annotations; columns represent predicted tracks: H3K4me1, H3K27me3, H3K27ac, ATAC, DNase, CAGE, and PRO-cap. These candidates were selected to illustrate the predicted bivalent dormancy signature and are not expected to match the top-ranked loci in panel c on a one-to-one basis. The color scale indicates *Z* scored predicted signal intensity, and gray indicates tracks not scored for a given candidate. Co-enrichment of H3K4me1 and H3K27me3 alongside suppression of H3K27ac, chromatin-accessibility, and transcriptional-output tracks defines the bivalent dormancy signature of prioritized poised/polycomb candidates. **(g)** Candidate-level external H3K27me3 support after strict TSS/GC-matched background comparison. Each point represents one DREC candidate summarized across public H3K27me3 datasets as log_2_ candidate-versus-matched-control signal. Human candidates are shown in blue and mouse candidates in orange. Because chromatin-class labels are non-exclusive, a candidate may contribute to more than one class-stratified group. In the poised/polycomb-versus-other-DREC contrast, each candidate is counted once as described in **Materials and Methods**. For this panel, the poised-enhancer-like and polycomb-marked classes shown separately in Fig. 3e were combined into a single poised/polycomb group. Poised/polycomb candidates show the strongest H3K27me3 enrichment over matched controls in both human and mouse, whereas active/scaffold, closed/unmarked, and low-accessibility candidates do not show the same positive shift. Detailed species-specific validation panels and direct poised/polycomb-versus-other-DREC contrasts are shown in **Supplementary Figs. 3 and 4.**

Of the 181 axolotl prior genes, 115 yielded primary resolvable mammalian ortholog assignments and were used for cross-species projection. Motif scanning of ortholog-centered genomic windows identified 202 human and 456 mouse motif-positive pre-candidate loci (**Supplementary Table 2**). After removing low-information motif clusters before AlphaGenome-based scoring, 199 human and 450 mouse intervals remained, comprising a 649-locus dual-species DREC atlas (**Fig. 3b**; **Supplementary Table 3**). The larger number of mouse pre-candidates likely reflects species-specific differences in motif-hit density, local gene-window architecture, genome annotation, and filtering behavior under the same motif-scanning criteria. The breadth of this atlas demonstrates that the axolotl-derived regenerative grammar is not an evolutionary relic restricted to a handful of conserved loci but is systematically represented across the mammalian genome within the syntenic neighborhoods of regeneration-associated genes.

To prioritize candidates most likely to represent dormant regulatory elements, we computed a DREC dormancy score (**Materials and Methods**) for each locus, integrating predicted H3K4me1^43^ and H3K27me3^44–46^ as positive contributors to a dormant/poised enhancer state and predicted H3K27ac^47^, chromatin accessibility^48,49^, and transcriptional output^50,51^ as negative contributors. This scoring framework was designed to distinguish between two fundamentally different fates that regenerative grammar could occupy in the mammalian genome: active deployment, in which motif-containing loci would show H3K27ac enrichment and chromatin accessibility, and epigenetic sequestration, in which the same grammar would be held under polycomb-mediated repression marked by H3K27me3 in the absence of active enhancer signatures^44–46^. Ranking by this score identified high-scoring candidates in both species (**Fig. 3c**), and top score bins were enriched for loci annotated as poised-enhancer-like and/or polycomb-marked relative to the full DREC atlas (**Fig. 3d**), whose broader non-exclusive chromatin-state composition is shown in **Fig. 3e**. Heatmap visualization of AlphaGenome-predicted epigenomic outputs across prioritized poised/polycomb candidates showed elevated H3K4me1 and H3K27me3 together with comparatively low H3K27ac, accessibility, and transcription-associated outputs (**Fig. 3f**), defining a bivalent dormancy signature analogous to that described at poised enhancers in embryonic and lineage-intermediate progenitor contexts^45,52^. Taken together, these findings indicate that the axolotl-derived regenerative grammar is not absent from mammalian genomes but is instead systematically preserved within orthologous gene neighborhoods while being predicted to be held in a state of epigenetic sequestration, providing the first genome-wide computational nomination of loci at which the regenerative deficit of adult mammals may reflect regulatory lockdown rather than genetic erasure.

### Poised/polycomb DRECs carry measured signatures of epigenetic dormancy and undergo postnatal attenuation of active-enhancer signal

The genome-wide identification of predicted bivalent dormancy at DREC loci raises a profound question at the intersection of evolutionary biology and regenerative medicine. Whether the mammalian lineage has genuinely lost the regulatory permission to execute a regenerative response, or instead retains a latent regenerative blueprint held under active epigenetic lockdown that could in principle be unlocked, cannot be resolved by computational prediction alone. We therefore compared the predicted bivalent dormancy signatures with three external support layers: measured epigenomic signal, developmental chromatin dynamics during the postnatal window in which regenerative capacity declines, and genomic overlap with human disease-associated variant annotations.

Candidate-level H3K27me3 signal, measured across public ENCODE/Roadmap epigenomic datasets^53–55^ and normalized against strict background intervals matched for chromosome, interval length, TSS distance, and GC content, was significantly elevated in poised/polycomb candidates relative to matched controls in both human and mouse (one-sided Wilcoxon signed-rank *P* = 4.48 × 10^−6^ and *P* = 3.82 × 10^−9^, respectively; **Fig. 3g**), and was higher than other DRECs in both species (Mann-Whitney U, human *P* = 1.68 × 10^−8^, mouse *P* = 9.70 × 10^−13^). Because predicted H3K27me3 contributes to the primary DREC dormancy score, we interpret measured H3K27me3 enrichment as external epigenomic support for the predicted chromatin state rather than as a feature-blind validation. The effect was consistent in direction and magnitude across species, with mouse poised/polycomb candidates showing a modestly stronger enrichment than their human counterparts. Other predicted classes, including active-scaffold, closed-unmarked, and low-accessibility candidates, showed weaker or less consistent H3K27me3 support, indicating that the signal was concentrated in the poised/polycomb subset rather than reflecting a uniform property of all DREC loci. Because these comparisons used strict matched backgrounds controlled for TSS distance and GC content, this pattern is less likely to be explained solely by gene proximity or GC content. Stratification of candidate-level H3K27me3 support by predicted chromatin class and direct contrast of poised/polycomb candidates against other DRECs confirmed that the enrichment signal is concentrated in the predicted poised/polycomb subset rather than distributed uniformly across the atlas in both mouse and human (**Supplementary Fig. 3a, b**; **Supplementary Fig. 4a, b**).

The second line of evidence addressed whether this epigenetic state is dynamically regulated during mammalian development rather than constitutively present throughout life. Mouse DRECs in the top 25% of DREC dormancy score that also carried poised/polycomb annotations showed postnatal attenuation of H3K27ac between postnatal day 1 and day 8: 38 of 48 loci decreased, with a median P8−P1 H3K27ac delta of −0.25 (one-sided Wilcoxon signed-rank *P* = 2.49 × 10^−4^; **Supplementary Fig. 3c**). This P1-to-P8 interval coincides with the decline of neonatal mouse cardiac regenerative capacity^8,9^, supporting postnatal attenuation of active-enhancer signal in prioritized poised/polycomb mDRECs.

The third line of evidence asked whether human DREC candidates overlap disease-associated variant annotations. Human DREC candidates showed increased overlap with a composite cardiac/muscle/neurologic GWAS Catalog trait set (25 of 199 candidates, 12.6%, versus 1,346 of 19,900 background intervals, 6.8%; odds ratio = 1.98; Fisher’s exact test, nominal *P* = 2.30 × 10^−3^; Bonferroni-adjusted *P* = 9.19 × 10^−3^). Cardiac overlap was nominally increased (16 of 199 candidates versus 958 of 19,900 backgrounds; odds ratio = 1.73; Fisher’s exact test, nominal *P* = 3.30 × 10^−2^) but did not remain significant after correction across the four predefined trait sets (adjusted *P* = 0.132). Neurologic and muscle overlaps were not significant (**Supplementary Fig. 4c**). **Supplementary Table 4** consolidates assay-level AlphaGenome-to-measured epigenome concordance, candidate-versus-background signal differences, GWAS overlap statistics, and phyloP/phastCons conservation metrics, providing a quantitative record of the validation and sensitivity analyses supporting the atlas.

Taken together, these analyses establish that AlphaGenome-prioritized poised/polycomb DRECs are not computational artifacts but represent a biologically coherent class of mammalian loci whose predicted dormancy is corroborated by measured epigenomic data, developmental chromatin dynamics, and human disease-variant architecture, supporting the model that the adult mammalian genome retains a latent regenerative blueprint held under active and measurable epigenetic control.

### *In silico* motif perturbation reveals that dormancy is actively enforced by sequence-encoded molecular locks conserved across mammalian evolution

Having established that bivalent dormancy at DREC loci is empirically supported and developmentally regulated, we asked whether this state is passively inherited from the surrounding chromatin environment or actively maintained by specific sequence elements encoded within the loci themselves. In this framework, genomic sequence and epigenomic state are treated as coupled layers. Sequence grammar defines the regulatory potential of a locus, whereas chromatin state determines whether that potential is deployed or repressed. Motif perturbation was therefore used to nominate sequence features that may stabilize dormancy. This distinction carries direct therapeutic implications: if dormancy is environmentally imposed, reactivation would require broad epigenetic reprogramming of the chromatin landscape, whereas if it is sequence-encoded, the targeted disruption of specific motif instances might be sufficient to shift individual loci toward a regeneration-competent state. To test this, we performed *in silico* motif perturbation at human-mouse matched orthologous DREC loci (**Fig. 1b**), reasoning that loci retained in both species would represent the most stringent and experimentally actionable test of dormant regenerative grammar. We focused on 18 matched human and mouse candidate loci corresponding to nine orthologous gene neighborhoods, spanning a range of predicted DREC dormancy scores and baseline chromatin states (**Supplementary Fig. 5**). In this primary human-mouse common ISM set, 78,931 native motif targets were evaluated in triplicate, yielding 236,793 replicate-level AlphaGenome perturbation effect estimates. This enabled locus-wide assessment of which native motifs contribute to the predicted dormant chromatin balance (**Fig. 4a**; **Materials and Methods**).

**Figure 4.**
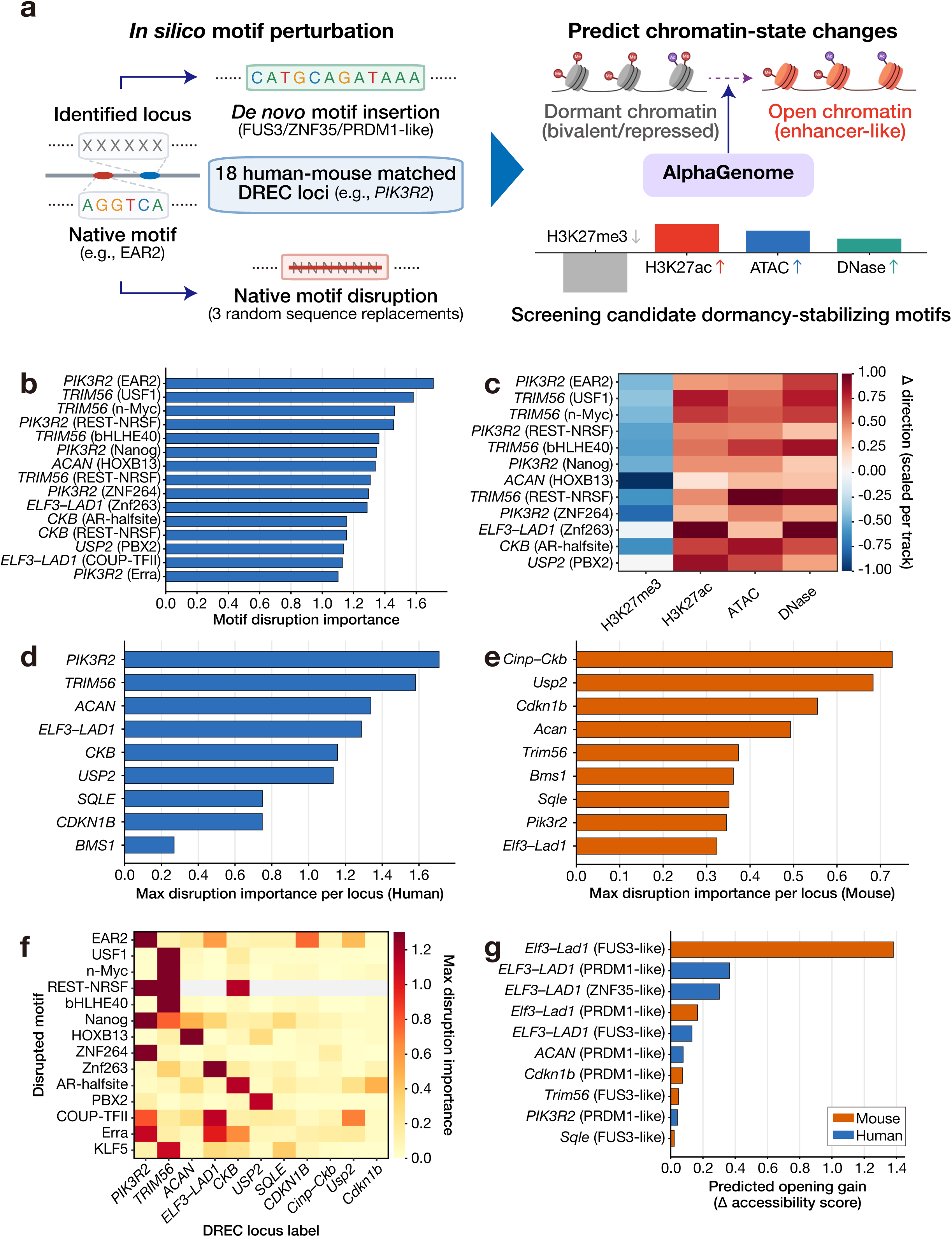
*In silico* motif perturbation identifies regulatory grammar components that maintain and can destabilize dormancy at human-mouse matched DREC loci. **(a)** Overview of the *in silico* motif-perturbation strategy applied to 18 human-mouse matched ortholog-associated DREC candidates. Native motif instances were disrupted by replacement with randomized sequences in three independent replicates, or the three selected *de novo* motifs (denovo_035, denovo_026 and denovo_033) were inserted at the candidate midpoint by same-length sequence replacement. The sequence illustrated in the schematic is denovo_035 (CATGCAGATAAA; closest known match, FUS3). AlphaGenome was used to predict the resulting changes in H3K27me3, H3K27ac, ATAC, and DNase signals, with disruption importance defined as the magnitude of predicted shift away from the dormant bivalent state and opening gain defined as the predicted shift toward an accessible, enhancer-like chromatin configuration. **(b)** Motif disruption importance scores across the top-ranked human-mouse matched DREC loci, all of which occur at human loci. Each bar represents the strongest disruption for a non-redundant locus-motif-family combination ranked by predicted disruption importance, reflecting the magnitude of predicted chromatin remodeling upon native motif disruption. The strongest effects are observed at *PIK3R2* (EAR2) and *TRIM56* (USF1, n-Myc, bHLHE40, and REST-NRSF) loci. **(c)** Directionality of predicted epigenomic changes upon native motif disruption at top-ranked loci, all of which occur at human loci. ISM was performed across the configured AlphaGenome output set where available, whereas this directionality heatmap displays H3K27me3, H3K27ac, ATAC, and DNase because these outputs most directly capture loss of repressive/poised signal and gain of enhancer/accessibility signal. The heatmap shows the predicted change relative to wild-type (variant minus WT, averaged over the central 4-kb summary window) for each displayed track, scaled independently per track to emphasize direction; note that absolute magnitudes differ by assay. Red indicates a predicted increase, blue a decrease. Disruption of these dormancy-stabilizing motifs is associated with H3K27me3 loss together with H3K27ac, ATAC, and DNase gain, consistent with dormancy destabilization. The heatmap shows 12 of the 15 locus-motif-family combinations displayed in Fig. 4b; three combinations — CKB (REST-NRSF), ELF3–LAD1 (COUP-TFII), and PIK3R2 (Erra) — lacked complete coverage across all four displayed tracks and were therefore omitted. **(d)** Locus-level summary of maximum disruption importance scores across human DREC loci. Each bar represents the maximum predicted disruption importance among all motif instances at the indicated locus; higher values indicate greater predicted sensitivity to motif disruption. *PIK3R2*, *TRIM56*, *ACAN*, and *ELF3–LAD1* show the strongest predicted responses. **(e)** Locus-level summary of maximum disruption importance scores across mouse DREC loci. The same metric as in (d) is shown for mouse orthologous loci. *Cinp–Ckb* and *Usp2* show the strongest predicted responses, with *Cdkn1b*, *Acan*, *Trim56*, and *Bms1* also displaying notable perturbation sensitivity. **(f)** Heatmap of maximum disruption importance scores stratified by disrupted motif family and DREC locus across the displayed 11-locus subset of the 18-locus human-mouse matched candidate set. The non-uniform distribution of high-importance values indicates that perturbation sensitivity reflects specific gene-motif combinations rather than overall motif density. Gray cells indicate gene-motif combinations in which the motif family was not detected at the locus. Loci are labeled by their nearest ortholog gene name, which denotes proximity within the search window rather than a validated target gene. **(g)** Predicted opening gain after *de novo* motif insertion for top-ranked human (blue) and mouse (orange) insertion events. Opening gain (Δ accessibility score) is the predicted shift toward an open, active chromatin state, defined as gains in H3K27ac, ATAC and DNase together with loss of H3K27me3, following insertion of a FUS3-like, PRDM1-like, or ZNF35-like *de novo* motif at the indicated locus. The strongest insertion response is observed at the mouse *Elf3–Lad1* locus (FUS3-like). Smaller positive effects are observed at human *ELF3–LAD1* and selected additional loci, indicating that *de novo* motif insertion can modulate dormant mammalian candidates but that effects are context-dependent. The overall more modest effects of *de novo* insertion relative to native motif disruption suggest that active repression rather than the absence of activating grammar is the dominant constraint at these loci.

The perturbation landscape revealed a finding that inverts the conventional understanding of transcription factor function at enhancers. The strongest motif disruptions did not simply reduce local enhancer potential, as would be expected if these motifs were activating elements. Instead, their computational ablation drove candidate loci away from the repressed bivalent state and toward an activated enhancer-like profile marked by loss of H3K27me3 together with gains in H3K27ac and chromatin accessibility (**Fig. 4b, c**; **Supplementary Fig. 6a, b**). This paradoxical result indicates that a subset of native motifs at DREC loci functions not to activate transcription but to actively enforce epigenetic repression, serving as candidate sequence features that may stabilize the dormant chromatin state. The strongest predicted effects were concentrated at the human *PIK3R2* and *TRIM56* loci, where disruption of motifs including EAR2, USF1, n-Myc, bHLHE40, and REST-NRSF produced coordinated H3K27me3 loss and enhancer activation signatures, suggesting that multiple dormancy-stabilizing motifs converge at individual loci to enforce repression through partially redundant mechanisms (**Fig. 4b, c; Supplementary Fig. 6b, c**). Perturbation sensitivity was observed across multiple loci in the analyzed 18-locus human-mouse matched set, but the mechanistic axis of the dormant-to-active transition was not uniform: at some loci, motif disruption primarily relieved H3K27me3-associated repression, whereas at others, the dominant response was gain of active enhancer marks with comparatively little change in H3K27me3, indicating that dormancy is maintained through locus-specific combinations of repressive mechanisms rather than a single shared pathway (**Fig. 4c–f**; **Supplementary Fig. 6d**).

To ask whether *de novo* motif insertion could increase predicted chromatin opening, we tested the same three prespecified eligible motifs described above—denovo_035, denovo_026 and denovo_033, with closest known matches to FUS3, ZNF35 and PRDM1, respectively—at the selected candidate loci. *De novo* motif insertion produced more modest and context-dependent effects than native motif disruption across most loci (**Fig. 4g**; **Supplementary Fig. 6e**), with the strongest response occurring at the mouse *Elf3–Lad1* locus (**Supplementary Fig. 6f**). The contrast between the potent activation-like shifts induced by native motif disruption and the limited reach of *de novo* insertion is not consistent with a model in which dormancy reflects the mere absence of activating grammar. Rather, it points to active sequence-encoded repression as the dominant constraint, implying that unlocking the latent regenerative potential at these loci will require the coordinated removal of dormancy-stabilizing elements rather than, or in addition to, the introduction of regenerative motifs.

The cross-species dimension of the perturbation results sharpens this conclusion into a broader evolutionary argument. The retention of perturbation-sensitive motif grammar within matched human and mouse DREC neighborhoods suggests that dormancy-stabilizing regulatory logic has been maintained during mammalian evolution^56,57^. The regenerative blueprint is therefore preserved in the mammalian genome not merely as a passive genomic fossil but as a sequence-encoded program held under active repressive control, in which specific native motif instances serve as the molecular locks that prevent its reactivation. The identity of these locks, their sequence-level conservation across species, and their predicted functional significance establish that the epigenetic silencing of regenerative grammar in adult mammals is not an irreversible evolutionary fait accompli but a regulatory state with a definable molecular architecture.

## Discussion

This study establishes an axolotl-to-mammal regulatory discovery framework for testing whether mammalian genomes have lost regenerative cis-regulatory information or retain it in an epigenetically locked state. Its novelty lies in coupling axolotl-derived regenerative motif grammar, projection through orthologous mammalian gene neighborhoods rather than direct enhancer-sequence alignment, AlphaGenome-based chromatin-state prediction, and *in silico* motif perturbation within a single framework^18–21^. By anchoring our discovery pipeline on axolotl limb regeneration chromatin-accessibility and single-cell transcriptomic data, we defined a combinatorial regenerative grammar enriched for HOX, nuclear receptor, SOX, AP-1, and additional injury- and patterning-associated transcription factor families. Projection of this grammar onto human and mouse orthologous gene neighborhoods, followed by AlphaGenome-based chromatin-state scoring, identified a 649-locus atlas of dormant regeneration enhancer candidates (DRECs). The central finding is that regeneration-associated regulatory grammar is not simply absent from mammalian genomes, but remains recoverable within selected orthologous neighborhoods where it is predicted to be epigenetically locked in a bivalent or polycomb-associated dormant state.

The predicted DREC state is characterized by co-enrichment of H3K4me1 and H3K27me3 with reduced H3K27ac, chromatin-accessibility signal, and transcriptional-output predictions^45–47^. Measured H3K27me3 enrichment in public human and mouse epigenomic datasets^53–55^, postnatal attenuation of H3K27ac at prioritized mouse DRECs during the P1-to-P8 interval^8,9^, and genomic overlap between human DRECs and disease-associated variant annotations support the biological coherence of this candidate class^58^. Together, these findings support a model in which loss of adult mammalian regenerative competence reflects regulatory sequestration of latent regenerative potential rather than erasure of the underlying cis-regulatory grammar.

The *in silico* motif perturbation analyses add a mechanistic layer to this model. Native motif disruption and *de novo* motif insertion address complementary questions: whether endogenous sequence features stabilize the predicted dormant state, and whether adding axolotl-derived regenerative motif input can increase predicted chromatin opening. Native motif disruption produced the stronger and more consistent shifts, moving selected loci away from a dormant/poised profile toward H3K27me3 loss with increased H3K27ac and accessibility-associated signals. Insertion of the three prespecified eligible *de novo* motifs produced smaller and more context-dependent effects, with positive opening gains at some loci and slight negative shifts at others. Together, these results argue against a simple model in which DREC dormancy reflects only the absence of activating regenerative motifs. Instead, they suggest that mammalian DREC loci contain native sequence features that function as candidate molecular locks, stabilizing dormancy while leaving opening potential accessible only in receptive sequence contexts or after release of endogenous constraints.

This framework also clarifies the relationship between genomic sequence and chromatin state. Sequence grammar defines the regulatory potential of a locus, whereas chromatin state determines whether that potential is deployed, poised, or repressed. In this view, mammalian DRECs represent regulatory neighborhoods in which axolotl-derived grammar, predicted bivalent dormancy, external epigenomic support, and sequence-perturbation sensitivity converge. Representative high-priority candidates, including *PIK3R2/Pik3r2*, *TRIM56/Trim56*, *ACAN/Acan*, *CKB/Cinp–Ckb*, and *ELF3–LAD1/Elf3–Lad1* neighborhoods, provide tractable entry points for massively parallel reporter assays (MPRA), locus-specific chromatin profiling, and CRISPR or base-editing perturbation^59–62^ of prioritized motif instances. This model reframes the postnatal loss of regenerative competence as a withdrawal of regulatory permission from loci that retain latent sequence potential but become constrained by polycomb-associated or low-accessibility chromatin.

Several limitations remain. The DREC atlas remains a computational prioritization resource and will require validation in injury-relevant mammalian cells and tissues. Future cell-type-resolved axolotl regulatory maps^14,15^ may refine the grammar assigned to blastema, epidermal, immune, nerve-associated, and stromal compartments. Because the single-cell-derived transcriptomic component contained one library per time point, pseudobulk expression results were used as expression-support evidence for prioritization rather than as definitive replicate-supported differential-expression claims. Finally, conservation of motif grammar does not establish conserved function, and linked gene labels do not prove target-gene causality. Despite these limitations, the DREC framework reframes mammalian regenerative failure as a testable problem of regulatory deployment. In this model, the mammalian genome has not simply lost the regulatory blueprint for regeneration; rather, selected components of that blueprint remain encoded within orthologous regulatory neighborhoods but are constrained by sequence-context and chromatin mechanisms that limit their activation. Whether perturbing these candidate molecular locks can restore regenerative chromatin states in living tissues remains to be tested, but the present work nominates the loci, motifs, and chromatin states needed to make that question experimentally addressable. DREC candidate intervals are motif-positive intervals generated by merging motif hits within ortholog-centered search windows rather than the windows themselves; they are large (median 517 kb in human and 166 kb in mouse) and should be read as regulatory neighborhoods rather than single enhancers; chromatin-state scores were nonetheless computed over a fixed 4,096-bp window centered on each candidate midpoint.

## Materials and Methods

### Study design and computational framework

We developed a regeneration-informed computational framework to identify dormant regeneration enhancer candidates (DRECs) in mammalian genomes. The framework was designed to capture the cis-regulatory logic of productive appendage regeneration in the axolotl and to transfer this information to orthologous regions in the human (hg38) and mouse (mm10) genomes. Stage-resolved axolotl limb-regeneration ATAC-seq and single-cell RNA-seq (scRNA-seq) data were first integrated to identify genes whose transcriptional changes were associated with dynamic chromatin accessibility during the established mid-bud regenerative phase. Regeneration-associated accessible regions were subsequently summarized at the level of transcription-factor (TF) motif families to define a transferable cis-regulatory grammar. The resulting gene and motif priors were used for the cross-species projection, AlphaGenome-based scoring, and *in silico* perturbation analyses described in **Figs. 3 and 4** and **Supplementary Figs. 5 and 6**.

### Axolotl limb-regeneration datasets

Bulk ATAC-seq data were obtained from NCBI BioProject PRJNA682840^24^ and comprised three biological replicates at each stage, except the 3-h stage, for which replicate 1 was excluded owing to low correlation as in the original study, leaving two replicates: homeostatic tissue at 0 h, acute trauma at 3 h, wound healing at 1 day, early bud at 3 days, mid-bud at 7 days, late bud at 14 days, palette at 22 days, and redifferentiated tissue at 33 days. The corresponding Sequence Read Archive accessions were SRR13268860–SRR13268883. Single-cell RNA-seq data were obtained from NCBI BioProject PRJNA589484^23^ (SRR10445716–SRR10445723), comprising one 10x Genomics Chromium library per regeneration stage. All analyses used the chromosome-scale UKY_AmexF1_1 axolotl genome assembly (RefSeq GCF_040938575.1; NCBI Assembly)^17,63^ and its corresponding NCBI RefSeq gene annotation^64^.

### ATAC-seq processing and peak identification

Paired-end ATAC-seq reads were aligned to the UKY_AmexF1_1 reference genome using BWA-MEM^65^. Alignments were processed with SAMtools^66^, and properly paired, uniquely mapped, non-duplicate fragments with a mapping quality of at least 30 were retained. Given the large size of the axolotl chromosomes, alignments were indexed using the CSI format.

To define stage-specific accessible chromatin landscapes, filtered alignments from the three biological replicates at each time point were merged and subjected to peak calling with MACS2^67^ in paired-end mode, using an effective genome size of 3.2 × 10^10^ bp and a false-discovery threshold of 0.05. Peaks were called independently for each of the eight stages. In parallel, alignments from all seven post-injury stages were pooled and subjected to peak calling under identical settings to generate a master set of 67,489 non-redundant accessible regions, which served as the reference coordinate system for all downstream comparisons. Stage-level accessibility tracks were generated from the merged alignments using deepTools^68^ and normalized to counts per million mapped fragments. For each master peak, accessibility was summarized both across the full peak interval and within a 500-bp window centered on the peak summit.

### Classification of regeneration-associated accessible regions

Peaks identified across the eight stages were merged by genomic overlap and annotated according to the stages in which they were detected. Dynamic regeneration-associated peaks were defined from stage-level peak-detection patterns after merging biological replicates within each time point. Specifically, regions absent at 0 h but detected in at least two post-injury stages were classified as dynamic regeneration-associated peaks. Requiring detection at more than one injury stage reduced the contribution of isolated peak calls and stage-specific technical variation. This catalog was used as an accessibility prior for downstream integration and cross-species projection, not as a formal replicate-level differential-accessibility call set.

Regions detected at 0 h were classified as constitutively accessible. Day-7-specific peaks were assigned to a single peak_annotation_class relative to RefSeq gene models using a mutually exclusive hierarchical scheme. Peaks within ±2 kb of a transcription start site (TSS) were classified as promoter-associated. Among the remaining peaks, those overlapping gene bodies but not annotated exons were classified as intronic. All remaining peaks were classified by nearest-TSS distance as proximal if the nearest TSS was ≤10 kb, distal if the nearest TSS was ≤100 kb, and intergenic otherwise. When a peak satisfied multiple criteria, the first matching class in this hierarchy was assigned. Raw feature-overlap annotations, including exon-overlap flags, were tracked during peak annotation but were not used as independent **Fig. 2c** classes or downstream support-score classes.

### scRNA-seq quantification and preprocessing

Single-cell transcriptomes were quantified using the kallisto | bustools framework^69,70^. A transcriptome index and transcript-to-gene map were generated from the UKY_AmexF1_1 genome and RefSeq annotation. Reads from each library were processed using the 10x Chromium v2 chemistry configuration, followed by cell-barcode correction, UMI-aware sorting, and gene-level count quantification. Count matrices from the eight time points were combined into a single AnnData^71^ object, with cell barcodes prefixed by library identifier to ensure uniqueness.

Quality control and downstream analyses were performed using Scanpy^72^. Genes detected in fewer than three cells were excluded. Cells were retained when they contained at least 200 detected genes, at least 500 total counts, and fewer than 20% mitochondrial transcripts. After filtering, the dataset comprised 232,501 cells and 26,299 genes. Counts were normalized to 10,000 counts per cell, log-transformed, and the 2,000 most variable genes were selected for dimensionality reduction. Principal-component analysis was performed using 30 components, and a nearest-neighbor graph was constructed with 15 neighbors. Leiden clustering^73^ was applied at a resolution of 0.6, and UMAP^74^ was used to visualize the transcriptional landscape across regeneration stages. All stochastic operations were performed with a fixed random seed of 17.

### Pseudobulk differential-expression analysis

Single-cell-derived pseudobulk expression profiles were generated to summarize stage-associated transcriptional dynamics while avoiding the treatment of individual cells as independent replicates^75,76^. Raw UMI counts were aggregated within each combination of time point and Leiden cluster. Leiden clusters were used as transcriptional-state bins for pseudobulk aggregation, not as definitive cell-type annotations. Because the scRNA-seq dataset contained one 10x Genomics library per time point, cluster-by-stage aggregates were used to summarize within-stage expression structure and reduce cell-level noise.

Model-based expression-support tables were generated using the quasi-likelihood framework implemented in edgeR^77^. Lowly expressed genes were removed using edgeR::filterByExpr with the comparison time-point group factor before TMM normalization^78^ with edgeR::calcNormFactors. Dispersion estimation and quasi-likelihood model fitting were then performed in edgeR. Primary contrasts compared days 1, 3, and 7 against the 0-h reference; additional contrasts compared day 7 with day 3 and day 14 with day 7. Model-derived *P* values were adjusted using the Benjamini-Hochberg procedure^79^. Genes with BH-adjusted *P* values below 0.05 were retained as FDR-thresholded expression-support genes for regeneration-prior construction and ranking, rather than being interpreted as definitive replicate-supported differentially expressed genes.

For visualization of temporal expression dynamics, pseudobulk counts were converted to log_2_ counts per million (log_2_CPM) and averaged within each time point. Expression values were standardized independently for each gene across the eight stages, and genes were hierarchically clustered using Euclidean distance with Ward linkage.

### Integration of chromatin accessibility and transcription

ATAC-seq and scRNA-seq data were integrated at the level of peak-gene associations to prioritize accessible regions linked to regeneration-associated transcriptional changes. Each master peak was assigned to genes whose TSS was located within 100 kb of the peak midpoint; when multiple genes fell within this interval, the gene with the nearest TSS was designated the primary assignment. For each peak-gene pair, an accessibility trajectory and an expression trajectory were constructed across the eight stages, and concordance was evaluated using Pearson correlation, including synchronous, temporally shifted, and overall trajectory comparisons. Contrast-specific ATAC-seq and RNA-seq fold changes, statistical significance, and directionality were also recorded.

A peak-gene pair was retained for a given contrast when the associated gene passed the BH-adjusted *P* < 0.05 threshold in the model-based pseudobulk expression-support table. These retained pairs are referred to as proximity-assigned peak-gene pairs retained through the FDR-thresholded expression-support filter and were used to construct the axolotl regeneration prior. The peak-gene assignments themselves were not subjected to a link-level hypothesis test or FDR correction. This filtering retained 1,364, 1,024, and 565 peak-gene pairs at days 1, 3, and 7 relative to 0 h, respectively. The day-7-versus-0-h comparison was used to define the regenerative prior for downstream mammalian analyses. Although earlier time points yielded more associations, they were expected to capture broad trauma, inflammatory, and wound-response programs. Day 7 corresponds to the established mid-bud blastema^14,24^ and therefore provides a more selective representation of the regulatory program supporting regenerative outgrowth. The 565 day-7-associated peak-gene pairs corresponded to 181 unique genes. An auxiliary support score (range 0–6), incorporating ATAC-seq and RNA-seq effect-size magnitude, accessibility-expression correlation, concordant directionality between modalities, peak annotation class, and RNA differential-expression significance, was calculated solely for candidate ranking and was not used to define statistical significance.

### Transcription-factor motif enrichment

Motif enrichment was performed to represent the axolotl regeneration program as a transferable regulatory grammar that could be projected across species despite limited conservation of individual enhancer sequences. The foreground comprised 417 unique accessible regions derived from significant day-7-versus-0-h peak-gene pairs that showed increased ATAC-seq accessibility. Each region was analyzed as a 301-bp sequence centered on the peak summit. Known and *de novo* motif enrichment was assessed using HOMER^80^ v5.1. Known-motif analysis used the vertebrate TF motif collection, and *de novo* motifs were assigned putative TF identities according to their closest known motif matches. The background comprised the remaining master ATAC-seq peaks, enabling enrichment to be assessed relative to accessible chromatin in general rather than randomly sampled genomic sequence.

Enriched motifs were also summarized into 13 curated TF families associated with developmental patterning, injury responses, and regenerative signaling: HOX, nuclear receptor, SOX, AP-1, KLF, TEAD, homeobox, WNT effectors, SMAD, bHLH, STAT, ETS, and NF-κB-related families^25–38^. Each family was summarized by the number of assigned enriched motifs and the strongest enrichment signal among its members, reported from the HOMER enrichment statistics as log-transformed *P* value support. Given the inherently combinatorial and partially redundant nature of TF grammar in accessible chromatin, these family-level summaries were used to prioritize and interpret motif classes rather than to define a statistically exhaustive catalog of regeneration-specific motifs. For cross-species projection, the axolotl-derived regulatory grammar consisted of the 181 regeneration-prior genes together with the scan-enabled enriched known and *de novo* motif set; the curated TF-family repertoire provided a manuscript-facing summary of this motif prior. Stage-level ATAC summaries, ATAC-RNA integration counts, motif-family enrichment summaries and *de novo* motif records are reported in **Supplementary Table 1**.

### Ortholog mapping and cross-species search windows

Axolotl regeneration-prior genes were mapped to human and mouse primary orthologs using a precomputed NCBI Datasets-derived ortholog table containing axolotl, human and mouse gene symbols. The ortholog table was normalized to standard axolotl, human and mouse symbol fields, and rows were classified as primary, ambiguous or unresolved. When orthology-type and confidence fields were present, one-to-one mappings and higher-confidence complete human-mouse assignments were prioritized; otherwise, unique complete symbol mappings were retained. Ambiguous multi-row assignments were flagged separately, and unresolved genes were excluded from cross-species projection. Of 181 axolotl prior genes, 115 primary ortholog rows were taken forward.

Human analyses used hg38 with GENCODE v48 primary-assembly gene annotation, and mouse analyses used mm10 with GENCODE vM23 gene annotation^81^. For each retained ortholog, the canonical gene span in the species-specific GTF was used to define a strand-aware transcription start site. Ortholog-centered search windows were generated using a 500 kb half-window around the target gene transcription start site. Overlapping gene-centered windows were merged before motif scanning.

Within each merged ortholog-centered search window, scan-enabled known and *de novo* motifs derived from the 7d versus 0h axolotl motif-enrichment analysis were scanned using HOMER ‘annotatePeaks.pl’ with ‘-size given’ and ‘-mbed’ output. Known motifs were selected from the HOMER vertebrate known-TF motif database using motif names recovered in the axolotl enrichment output, and *de novo* motifs were copied from the primary HOMER *de novo* motif files. The scan used the full selected known/*de novo* motif set; the 13 curated TF-family groups were used as a family-level summary of this motif prior rather than as the complete set of scanned motifs.

Motif hits were assigned back to overlapping ortholog-centered windows and clustered into candidate loci by merging hits separated by no more than 128 bp. Candidate clusters were summarized by total motif-hit count, number of unique motifs, known and *de novo* hit counts, assigned regeneration TF families, linked mammalian genes, linked axolotl prior genes, motif-source enrichment strength, and distance from the linked target-gene TSS. Candidate loci were ordered by decreasing unique motif count, decreasing total motif-hit count, decreasing motif-source enrichment strength, and increasing distance to the linked target-gene TSS. Ortholog mappings, motif-positive pre-candidates, the nine-candidate exclusion audit, chromatin-class counts, score-bin summaries and top-ranked candidate audits are reported in **Supplementary Table 2**.

### DREC candidate catalog construction

Candidate preselection for AlphaGenome scoring required at least two motif hits and at least two unique motifs. Candidate loci passing these criteria were carried forward for AlphaGenome scoring and final catalog construction. Pre-candidates not retained for AlphaGenome scoring were audited by exact candidate identifier matching. Excluded loci were low-information short clusters with insufficient motif support. Candidate construction and filtering are reported in **Supplementary Table 2**, and the final DREC candidate universe is reported in **Supplementary Table 3**.

### AlphaGenome chromatin-state scoring

DREC candidates were scored using a local AlphaGenome implementation. For each candidate, a 131,072-bp input sequence window was extracted from the corresponding reference genome. Predicted regulatory outputs were summarized over a central 4,096-bp window at 128-bp resolution. Additional flanking and plotting windows were set to ±10 kb. Predicted track groups included H3K4me1, H3K27me3, H3K27ac, ATAC, DNase, CAGE, and PRO-cap where available. Center and flank signal summaries were computed as mean predicted signal values.

For tracks available in both species, center-window signal values were converted to track-specific *Z* scores across the combined set of 649 scored human and mouse candidates. PRO-cap, which was available only for human candidates, was standardized across the 199 human candidates. For candidate *i* and track *t*, the *Z* score was defined as:

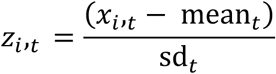

where *x*_i,t_ is the center-window signal for candidate *i*, and *mean_t_* and *sd_t_* are the mean and standard deviation of the same track across the combined candidate set, except for human-only PRO-cap.

The bivalency core score was defined as:

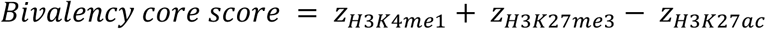

The dormancy support score was defined as:

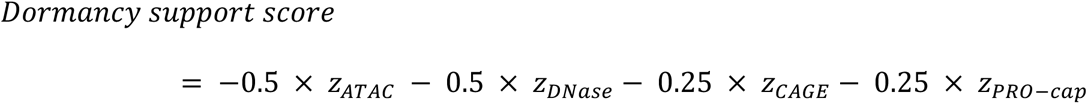

The final DREC dormancy score was defined as:

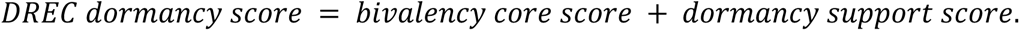

Thus, H3K4me1 and H3K27me3 contributed positively to the score, whereas H3K27ac, chromatin-accessibility tracks, and transcriptional-output tracks contributed negatively. If a predicted output was unavailable for a candidate, the corresponding term was treated as neutral and contributed 0 to the composite score. Because PRO-cap predictions were unavailable for all 450 mouse candidates, the number of contributing terms differs between species. DREC dormancy scores were computed from the track-specific *Z* scores described above. The composite score itself was not re-standardized and was ranked within species. Because most tracks were standardized across the combined human and mouse candidate set, DREC dormancy scores are broadly comparable across species; they differ only by the human-specific PRO-cap contribution, which should be kept in mind when interpreting the combined ranking in **Fig. 3c**. AlphaGenome outputs were also used to assign predicted chromatin-state labels and to prioritize dormant/poised-like candidate intervals. For representative locus-style visualizations, selected candidates were centered on the DREC interval and displayed as schematic H3K4me1, H3K27me3, H3K27ac, and accessibility profiles scaled from centered AlphaGenome summary values.

### Chromatin class assignment

Candidate chromatin labels were assigned from AlphaGenome-derived predicted *Z* score features. Labels were non-exclusive; a candidate could carry more than one class.

Poised-enhancer-like candidates were defined as loci satisfying:

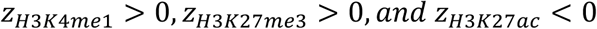

Polycomb-marked candidates were defined as loci satisfying:

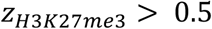

Closed-unmarked candidates were defined as loci satisfying:

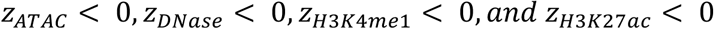

Active-scaffold candidates were defined as loci satisfying:

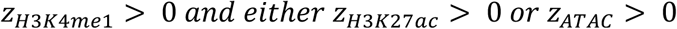

Low-accessibility candidates were defined as loci satisfying:

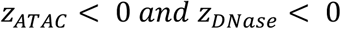

For figure summaries, semicolon-delimited class labels were expanded into long format before counting by species and chromatin class. Because labels were non-exclusive, class-count totals could exceed the number of unique loci. Poised-enhancer-like and polycomb-marked labels defined the dormant/poised-like chromatin-state subsets used for stratified validation analyses, whereas active-scaffold, closed-unmarked, and low-accessibility labels captured additional candidate-state structure. Candidates that satisfied none of the five chromatin-class definitions were labeled unclassified (n = 13; 3 human and 10 mouse). These candidates were retained in **Supplementary Table 3** but were not included in the five-class count panel in **Fig. 3e**. Because the *Z* scores were defined relative to the analyzed candidate population, these chromatin-class labels are operational prioritization categories rather than absolute tissue- or cell-type-specific chromatin-state assignments.

### Matched backgrounds and centered validation intervals

Matched genomic backgrounds were generated to compare DREC candidates with random genomic intervals of similar genomic context. Each candidate was assigned 100 chromosome- and length-matched random background intervals, excluding blacklist-overlapping regions where applicable. Background-interval counts and failed-match audits are summarized in **Supplementary Table 4** where applicable. External validation analyses were performed using centered 512-bp intervals generated from candidate midpoints. Centered intervals standardized scoring width across candidates and provided a consistent sequence window for downstream assay-design resources. To assess sensitivity to genomic confounding, an additional strict background set was generated with matching for transcription start site distance and GC content. Strict backgrounds were used for sensitivity analyses of epigenomic enrichment and conservation comparisons. Candidate intervals for which strict matches could not be generated were excluded from strict-background comparisons rather than replaced by unmatched intervals.

### Public epigenomic signal scoring

Public epigenomic signal was scored over centered 512-bp candidate and background intervals. Human analyses included ENCODE/Roadmap-like H3K27ac (7 tracks) and H3K27me3 (6 tracks); no interval-compatible human H3K4me1 track was available, and the corresponding term was therefore treated as neutral for all human candidates. Mouse analyses included H3K4me1 (9 tracks), H3K27ac, and H3K27me3 across multiple tissues. Measured signal was summarized as mean signal over each centered interval. Candidate and background distributions were compared by species, tissue, assay, and background type. Candidate-vs-background comparisons were summarized using mean and median signal differences and Mann-Whitney U tests. Public epigenome support was summarized through candidate-level AlphaGenome-to-measured concordance and subset-specific measured support. Strict TSS- and GC-matched backgrounds were used to test candidate-versus-background signal after controlling for local genomic context.

For the main external-support panel and the human and mouse supplementary validation panels, public H3K27me3 support was summarized at the candidate level. For each DREC candidate, log_2_ candidate-versus-matched-control signal was summarized across available public H3K27me3 datasets, yielding one candidate-level support value per plotted point. Candidate-level H3K27me3 support panels included only DREC candidates with available public H3K27me3 signal after matched-background summarization, resulting in 196 of 199 hDRECs and 440 of 450 mDRECs in the plotted support analyses. Candidate chromatin-class labels were expanded in long format for class-stratified panels because labels were non-exclusive. For subset contrasts, each candidate was counted once, with the poised/polycomb label prioritized over other labels when a candidate carried multiple labels. Poised/polycomb candidates were compared with all other DRECs using a one-sided Mann-Whitney U test. Within-class candidate-level support against zero was evaluated using one-sided Wilcoxon signed-rank tests. Significance symbols in the figures use conventional thresholds (*\*P* < 0.05, *\*\*P* < 0.01, *\*\*\*P* < 0.001, *\*\*\*\*P* < 0.0001), while detailed values are summarized in **Supplementary Tables 1–5**.

### AlphaGenome-to-measured epigenome concordance

For each candidate, AlphaGenome-predicted values were paired with measured public epigenomic signal for corresponding assay classes. Concordance was quantified using Spearman rank correlation across candidate loci. Human and mouse concordance results were summarized by assay. Assay-level concordance summaries were retained as **Supplementary Table 4**. Candidate-level H3K27me3 support was used for the main external-validation figure and the human and mouse supplementary validation figures.

### Mouse P1/P8 histone and scATAC support

Mouse postnatal histone support was evaluated using public mouse ventricle H3K27ac and H3K27me3 tracks from GSE123867. Candidate and background centered intervals were scored over the available histone-mark tracks. For the supplementary postnatal analysis, we focused on active-enhancer signal attenuation at prioritized poised/polycomb mDREC loci by summarizing H3K27ac signal as P8−P1 deltas. The primary postnatal test evaluated whether P8−P1 H3K27ac deltas were below zero using a one-sided Wilcoxon signed-rank test. H3K27me3 signal from the same dataset was retained as a contextual histone-mark support layer, but the main postnatal claim was based on H3K27ac attenuation. Mouse injury/scATAC support was summarized using GSE153479-derived accessibility data and reported as a context-specific support layer.

### GWAS variant overlap

Trait sets were defined using regular-expression filters as a composite cardiac/muscle/neurologic category and separate cardiac, muscle and neurologic categories. Centered 512-bp candidate and background intervals were expanded by 10 kb on each side before overlap with GWAS Catalog annotations, and candidate and background overlap fractions were then compared using one-sided Fisher’s exact tests. Bonferroni correction was applied separately within each candidate subset across the four predefined trait sets. The trait-filter definitions used for the analysis are reported in **Supplementary Table 4**. GWAS Catalog associations were obtained from the NHGRI–EBI GWAS Catalog full associations download (gwas-catalog-download-associations-v1.0-full.tsv), downloaded on 21 May 2026. The analyzed snapshot was dated 17 May 2026, contained 1,124,004 association records, and included records added to the Catalog through 14 May 2026.

### Evolutionary conservation and grammar-conservation analysis

Evolutionary conservation was assessed using UCSC phyloP and phastCons tracks. Human analyses used 100-way conservation tracks, and mouse analyses used 60-way conservation tracks. Candidate and background intervals were scored using interval-level conservation summaries. Associations between motif grammar features and phyloP/phastCons conservation scores were evaluated using Spearman rank correlation separately for human and mouse. GERP++ was not included because a clean hg38/mm10 whole-genome interval-compatible track was not available in the current analysis environment. Conservation and grammar-conservation outputs are retained as **Supplementary Table 4**.

### *In silico* motif perturbation and *de novo* motif insertion

Motif-level *in silico* perturbation was performed for two predefined DREC subsets. The primary human-mouse common ISM set comprised 18 matched human and mouse DREC loci selected on the basis of high DREC dormancy score in both species and availability of a confident one-to-one ortholog pair. These loci corresponded to the following gene neighborhoods: *BMS1/Bms1*, *CKB/Cinp–Ckb*, *TRIM56/Trim56*, *CDKN1B/Cdkn1b*, *PIK3R2/Pik3r2*, *ACAN/Acan*, *USP2/Usp2*, *ELF3–LAD1/Elf3–Lad1*, and *SQLE/Sqle*. Per-locus DREC dormancy scores and baseline predicted epigenomic states of these loci are summarized in **Supplementary Fig. 5** and **Supplementary Table 5**.

For native motif disruption, all PWM-significant motif instances within a 16,384-bp window centered on the candidate locus midpoint at each locus were enumerated. Each native motif target was individually disrupted by replacing the motif sequence with a randomly generated sequence of identical length, sampled uniformly over A/C/G/T, and this procedure was repeated three times with independent random seeds. The disrupted sequence was substituted into the reference 131 kb AlphaGenome input window, and AlphaGenome was used to predict the resulting changes in H3K4me1, H3K27me3, H3K27ac, ATAC, DNase, CAGE, and PRO-cap signals within the 4,096-bp central scoring window relative to the wild-type prediction. For the human-mouse common 18-locus ISM set, 78,931 native motif targets were evaluated in triplicate, yielding 236,793 replicate-level motif-disruption effect estimates. An extended top-dormant 20-locus ISM set was analyzed in parallel as a broader prioritization resource. This set contained 89,720 native motif targets and yielded 269,160 triplicate disruption estimates. Results from the common 18-locus set and the top-dormant 20-locus set are reported as separate analysis sets in **Supplementary Table 5**.

Disruption importance for each motif instance was defined as the reduction, relative to the wild-type sequence, in a composite poised score (i.e., −Δ poised log- score). The poised score was computed as a weighted sum of log_2_(1 + central-window signal) across the seven predicted tracks, with positive weights for H3K4me1 (+1) and H3K27me3 (+1) and negative weights for H3K27ac (−1), ATAC (−0.5), DNase (−0.5), CAGE (−0.25) and PRO-cap (−0.25); higher disruption importance thus indicates a larger predicted shift away from the dormant poised state. Motif instances were ranked by disruption importance within each locus. For each native motif target, the maximum disruption importance across the three independent random-replacement replicates was retained; the maximum target-level value within each locus was used as the locus-level summary statistic. Motif-by-locus heatmaps were constructed from the maximum target-level disruption importance for each motif-family-locus combination. Prioritized motif-level ISM outputs, including per-locus summary statistics for the 18 matched loci—DREC dormancy score, maximum motif-disruption importance and its top-ranked motif, the predicted H3K27me3 and H3K27ac changes at that motif, and the maximum *de novo* insertion opening gain—are reported in **Supplementary Table 5**. The relationship between disruption importance and predicted opening gain, the H3K27me3-to-H3K27ac signature of the strongest disruptions, and per-locus maximum disruption importance are shown in **Supplementary Fig. 6a, b, and d**, respectively.

For *de novo* motif insertion, eligibility was determined before AlphaGenome outcome evaluation. Only primary HOMER *de novo* motif files matching the canonical motif <number>.motif pattern, with scan_enabled = TRUE and an unambiguous A/C/G/T consensus sequence, were considered eligible; auxiliary reverse-complement and similar-motif entries were excluded. Eligible motifs were ranked by decreasing HOMER-derived source_log_p_support, and the top three were selected without hard-coded motif IDs: denovo_035 (HOMER header rank 24; CATGCAGATAAA; closest known match, FUS3), denovo_026 (header rank 15; TAGAATTGTGAA; closest known match, ZNF35), and denovo_033 (header rank 22; GCACTGAGAAAG; closest known match, PRDM1). The source_log_p_support field is the positive HOMER-derived support statistic recorded in **Supplementary Table 1**; larger values indicate stronger support and the field should not be interpreted as a raw *P* value. Each selected motif was inserted at the candidate midpoint by replacing an equal-length segment of the reference sequence without altering flanking nucleotides. Closest-match labels are sequence-similarity annotations and do not establish activity of the named transcription factors. AlphaGenome was then used to predict the resulting opening gain, defined as the increase, relative to wild-type, in an accessibility score computed as a weighted sum of log_2_(1 + central-window signal). The opening gain score used positive weights for H3K27ac (+1), ATAC (+1), and DNase (+1), and a negative weight for H3K27me3 (−1), such that higher values indicate a predicted shift toward an accessible, enhancer-like chromatin state. Per-locus opening gains are shown in **Supplementary Fig. 6e**; representative wild-type-versus-variant prediction tracks for the strongest disruption (*PIK3R2*) and insertion (*Elf3–Lad1*) events are shown in **Supplementary Fig. 6c and f**, respectively.

### Statistical analysis and reproducibility

Single-cell-derived pseudobulk expression-support tables were generated using edgeR quasi-likelihood models with Benjamini-Hochberg correction. Because the scRNA-seq dataset contained one library per time point, BH-adjusted *P* < 0.05 entries were used as FDR-thresholded filtering and ranking support for regeneration-prior construction, not as standalone replicate-supported differential-expression calls. Accessibility-expression relationships were evaluated using Pearson correlation. Motif enrichment was assessed using HOMER enrichment *P* values.

ATAC-seq data were processed using BWA-MEM, SAMtools, MACS2, and deepTools. Single-cell data were quantified and analyzed using kallisto, bustools, Scanpy, and AnnData. Leiden clustering and UMAP were performed with fixed random seeds for reproducibility. Differential-expression analysis used edgeR. Motif enrichment and ortholog-centered motif scanning were performed using HOMER v5.1, including annotatePeaks.pl for cross-species search-window scanning. bigWig-derived signal extraction and interval-level public epigenomic signal summaries were performed using pyBigWig. Software tools and key analysis parameters are described in the relevant **Materials and Methods** sections.

Candidate-versus-background measured epigenomic signal was compared using Mann-Whitney U tests and summarized using candidate/background mean and median differences. Candidate-level H3K27me3 support in poised/polycomb DRECs was evaluated against zero using one-sided Wilcoxon signed-rank tests and compared with other DRECs using one-sided Mann-Whitney U tests. Mouse postnatal H3K27ac attenuation at prioritized poised/polycomb mDREC loci was evaluated using a one-sided Wilcoxon signed-rank test of P8−P1 H3K27ac deltas against zero, with the alternative hypothesis that the median delta was less than zero. AlphaGenome-to-measured epigenome concordance and grammar-conservation relationships were assessed using Spearman rank correlation. GWAS annotation-overlap enrichment was assessed using Fisher’s exact test with a greater-tail alternative. Multiple-testing adjusted significance was reported where provided by upstream differential-expression or motif-enrichment outputs; otherwise, exact test statistics and nominal *P* values are reported in the numbered Supplementary Tables where applicable.

## Data availability

Public datasets and genome references used in this study are identified in the **Materials and Methods** and **Supplementary Table 4**, with accession identifiers provided where applicable. Numbered **Supplementary Tables 1–5** are provided with this preprint. **Supplementary Table 1** contains stage-level axolotl ATAC summaries, ATAC-RNA integration counts, motif-family enrichment summaries and *de novo* motif records. **Supplementary Table 2** contains ortholog-mapping, candidate-construction and quality-control summaries. **Supplementary Tables 3–5** contain the final dual-species DREC catalog, external-support and sensitivity analyses, and locus-level *in silico* mutagenesis summaries, respectively. The GWAS Catalog input snapshot used in this study had SHA-256 checksum ea97dc7c060f7277ba1b974c878511a1e22a664ee96cadcdc58c490c83874118.

## Acknowledgements

This work was supported by KAKENHI grants from the Japan Society for the Promotion of Science (JSPS) to H.S. (23K28184, 25H01571, and 26H00766), JST FOREST Program to H.S. (JPMJFR242Q), as well as the Canon Foundation and Nakatani Foundation. Figures were created in part with BioRender.com. We thank T. Suzuoka, T. Sakuma, and T. Ito for critical reading of the manuscript, all laboratory members for helpful discussions, and K. Tanaka for assistance with manuscript preparation.

## Author Contributions

HS initially conceived of and supervised the project. TF and KN performed all formal analyses with the help of TS. TF, KN and TS prepared all display items, and HS mainly wrote the draft version of the manuscript with inputs from all authors. All authors approved the final manuscript.

## References

1. Poss, K. D. Advances in understanding tissue regenerative capacity and mechanisms in animals. Nat. Rev. Genet. 11, 710–722 (2010).

2. Brockes, J. P. & Kumar, A. Appendage regeneration in adult vertebrates and implications for regenerative medicine. Science 310, 1919–1923 (2005).

3. Tanaka, E. M. & Reddien, P. W. The cellular basis for animal regeneration. Dev. Cell 21, 172–185 (2011).

4. Poss, K. D. & Tanaka, E. M. Hallmarks of regeneration. Cell Stem Cell 31, 1244–1261 (2024).

5. Poss, K. D., Wilson, L. G. & Keating, M. T. Heart regeneration in zebrafish. Science 298, 2188–2190 (2002).

6. Gemberling, M., Bailey, T. J., Hyde, D. R. & Poss, K. D. The zebrafish as a model for complex tissue regeneration. Trends Genet. 29, 611–620 (2013).

7. Godwin, J. W., Pinto, A. R. & Rosenthal, N. A. Macrophages are required for adult salamander limb regeneration. Proc. Natl. Acad. Sci. U. S. A. 110, 9415–9420 (2013).

8. Porrello, E. R. et al. Transient regenerative potential of the neonatal mouse heart. Science 331, 1078–1080 (2011).

9. Wang, Z. et al. Mechanistic basis of neonatal heart regeneration revealed by transcriptome and histone modification profiling. Proc. Natl. Acad. Sci. U. S. A. 116, 18455–18465 (2019).

10. Kang, J. et al. Modulation of tissue repair by regeneration enhancer elements. Nature 532, 201–206 (2016).

11. Pfefferli, C. & Jaźwińska, A. The careg element reveals a common regulation of regeneration in the zebrafish myocardium and fin. Nat. Commun. 8, 15151 (2017).

12. Thompson, J. D. et al. Identification and requirements of enhancers that direct gene expression during zebrafish fin regeneration. Development 147, dev191262 (2020).

13. Harris, R. E. Regeneration enhancers: a field in development. Am. J. Physiol. Cell Physiol. 323, C1548–C1554 (2022).

14. Gerber, T. et al. Single-cell analysis uncovers convergence of cell identities during axolotl limb regeneration. Science 362, eaaq0681 (2018).

15. Leigh, N. D. et al. Transcriptomic landscape of the blastema niche in regenerating adult axolotl limbs at single-cell resolution. Nat. Commun. 9, 5153 (2018).

16. Nowoshilow, S. et al. The axolotl genome and the evolution of key tissue formation regulators. Nature 554, 50–55 (2018).

17. Smith, J. J. et al. A chromosome-scale assembly of the axolotl genome. Genome Res. 29, 317–324 (2019).

18. Long, H. K., Prescott, S. L. & Wysocka, J. Ever-changing landscapes: Transcriptional enhancers in development and evolution. Cell 167, 1170–1187 (2016).

19. Villar, D. et al. Enhancer evolution across 20 mammalian species. Cell 160, 554–566 (2015).

20. Visel, A. et al. Ultraconservation identifies a small subset of extremely constrained developmental enhancers. Nat. Genet. 40, 158–160 (2008).

21. Avsec, Ž., et al. Advancing regulatory variant effect prediction with AlphaGenome. Nature 649, 1206–1218 (2026).

22. Kumar, S. et al. TimeTree 5: An expanded resource for species divergence times. Mol. Biol. Evol. 39, msac174 (2022).

23. Li, H. et al. Dynamic cell transition and immune response landscapes of axolotl limb regeneration revealed by single-cell analysis. Protein Cell 12, 57–66 (2021).

24. Wei, X. et al. An ATAC-seq dataset uncovers the regulatory landscape during axolotl limb regeneration. Front. Cell Dev. Biol. 9, 651145 (2021).

25. Gardiner, D. M., Blumberg, B., Komine, Y. & Bryant, S. V. Regulation of HoxA expression in developing and regenerating axolotl limbs. Development 121, 1731–1741 (1995).

26. Sarkar, A. & Hochedlinger, K. The sox family of transcription factors: versatile regulators of stem and progenitor cell fate. Cell Stem Cell 12, 15–30 (2013).

27. Gardiner, D. M. & Bryant, S. V. Molecular mechanisms in the control of limb regeneration: the role of homeobox genes. Int. J. Dev. Biol. 40, 797–805 (1996).

28. Cadigan, K. M. & Waterman, M. L. TCF/LEFs and Wnt signaling in the nucleus. Cold Spring Harb. Perspect. Biol. 4, a007906 (2012).

29. Kawakami, Y. et al. Wnt/beta-catenin signaling regulates vertebrate limb regeneration. Genes Dev. 20, 3232–3237 (2006).

30. Beisaw, A. et al. AP-1 contributes to chromatin accessibility to promote sarcomere disassembly and cardiomyocyte protrusion during zebrafish heart regeneration. Circ. Res. 126, 1760–1778 (2020).

31. Herrera, S. C. & Bach, E. A. JAK/STAT signaling in stem cells and regeneration: from Drosophila to vertebrates. Development 146, dev167643 (2019).

32. Karin, M. & Clevers, H. Reparative inflammation takes charge of tissue regeneration. Nature 529, 307–315 (2016).

33. Denis, J. F. et al. Activation of Smad2 but not Smad3 is required to mediate TGF- β signaling during axolotl limb regeneration. Development 143, 3481–3490 (2016).

34. Nguyen, M. et al. Retinoic acid receptor regulation of epimorphic and homeostatic regeneration in the axolotl. Development 144, 601–611 (2017).

35. Bialkowska, A. B., Yang, V. W. & Mallipattu, S. K. Krüppel-like factors in mammalian stem cells and development. Development 144, 737–754 (2017).

36. Dey, A., Varelas, X. & Guan, K. L. Targeting the Hippo pathway in cancer, fibrosis, wound healing and regenerative medicine. Nat. Rev. Drug Discov. 19, 480–494 (2020).

37. Zammit, P. S. Function of the myogenic regulatory factors Myf5, MyoD, Myogenin and MRF4 in skeletal muscle, satellite cells and regenerative myogenesis. Semin. Cell Dev. Biol. 72, 19–32 (2017).

38. Remy, P. & Baltzinger, M. The Ets-transcription factor family in embryonic development: lessons from the amphibian and bird. Oncogene 19, 6417–6431 (2000).

39. Werling, U. & Schorle, H. Transcription factor gene AP-2 gamma essential for early murine development. Mol. Cell. Biol. 22, 3149–3156 (2002).

40. Takeuchi, T. et al. Newt Hoxa13 has an essential and predominant role in digit formation during development and regeneration. Development 149, dev200282 (2022).

41. Scott, R. W., Arostegui, M., Schweitzer, R., Rossi, F. M. V. & Underhill, T. M. Hic1 defines quiescent mesenchymal progenitor subpopulations with distinct functions and fates in skeletal muscle regeneration. Cell Stem Cell 25, 797–813.e9 (2019).

42. Thummel, R., Ju, M., Sarras, M. P. & Godwin, A. R. Both Hoxc13 orthologs are functionally important for zebrafish tail fin regeneration. Dev. Genes Evol. 217, 413–420 (2007).

43. Heintzman, N. D. et al. Distinct and predictive chromatin signatures of transcriptional promoters and enhancers in the human genome. Nat. Genet. 39, 311–318 (2007).

44. Margueron, R. & Reinberg, D. The Polycomb complex PRC2 and its mark in life. Nature 469, 343–349 (2011).

45. Rada-Iglesias, A. et al. A unique chromatin signature uncovers early developmental enhancers in humans. Nature 470, 279–283 (2011).

46. Bernstein, B. E. et al. A bivalent chromatin structure marks key developmental genes in embryonic stem cells. Cell 125, 315–326 (2006).

47. Creyghton, M. P. et al. Histone H3K27ac separates active from poised enhancers and predicts developmental state. Proc. Natl. Acad. Sci. U. S. A. 107, 21931–21936 (2010).

48. Thurman, R. E. et al. The accessible chromatin landscape of the human genome. Nature 489, 75–82 (2012).

49. Buenrostro, J. D. et al. Transposition of native chromatin for fast and sensitive epigenomic profiling of open chromatin, DNA-binding proteins and nucleosome position. Nat. Methods 10, 1213–1218 (2013).

50. Andersson, R. et al. An atlas of active enhancers across human cell types and tissues. Nature 507, 455–461 (2014).

51. Core, L. J. et al. Analysis of nascent RNA identifies a unified architecture of initiation regions at mammalian promoters and enhancers. Nat. Genet. 46, 1311–1320 (2014).

52. Wang, A. et al. Epigenetic priming of enhancers predicts developmental competence of hESC-derived endodermal lineage intermediates. Cell Stem Cell 16, 386–399 (2015).

53. ENCODE Project Consortium. An integrated encyclopedia of DNA elements in the human genome. Nature 489, 57–74 (2012).

54. Roadmap Epigenomics Consortium et al. Integrative analysis of 111 reference human epigenomes. Nature 518, 317–330 (2015).

55. ENCODE Project Consortium et al. Expanded encyclopaedias of DNA elements in the human and mouse genomes. Nature 583, 699–710 (2020).

56. Kumar, S. & Hedges, S. B. A molecular timescale for vertebrate evolution. Nature 392, 917–920 (1998).

57. Álvarez-Carretero, S. et al. A species-level timeline of mammal evolution integrating phylogenomic data. Nature 602, 263–267 (2022).

58. Cerezo, M. et al. The NHGRI-EBI GWAS Catalog: standards for reusability, sustainability and diversity. Nucleic Acids Res. 53, D998–D1005 (2025).

59. Jinek, M. et al. A programmable dual-RNA-guided DNA endonuclease in adaptive bacterial immunity. Science 337, 816–821 (2012).

60. Cong, L. et al. Multiplex genome engineering using CRISPR/Cas systems. Science 339, 819–823 (2013).

61. Mali, P. et al. RNA-guided human genome engineering via Cas9. Science 339, 823–826 (2013).

62. Komor, A. C., Kim, Y. B., Packer, M. S., Zuris, J. A. & Liu, D. R. Programmable editing of a target base in genomic DNA without double-stranded DNA cleavage. Nature 533, 420–424 (2016).

63. Schloissnig, S. et al. The giant axolotl genome uncovers the evolution, scaling, and transcriptional control of complex gene loci. Proc. Natl. Acad. Sci. U. S. A. 118, e2017176118 (2021).

64. O’Leary, N. A. et al. Exploring and retrieving sequence and metadata for species across the tree of life with NCBI Datasets. Sci. Data 11, 732 (2024).

65. Li, H. Aligning sequence reads, clone sequences and assembly contigs with BWA-MEM. arXiv:1303.3997 (2013).

66. Li, H. et al. The Sequence Alignment/Map format and SAMtools. Bioinformatics 25, 2078–2079 (2009).

67. Zhang, Y. et al. Model-based analysis of ChIP-Seq (MACS). Genome Biol. 9, R137 (2008).

68. Ramírez, F., Dündar, F., Diehl, S., Grüning, B. A. & Manke, T. deepTools: a flexible platform for exploring deep-sequencing data. Nucleic Acids Res. 42, W187–W191 (2014).

69. Bray, N. L., Pimentel, H., Melsted, P. & Pachter, L. Near-optimal probabilistic RNA-seq quantification. Nat. Biotechnol. 34, 525–527 (2016).

70. Melsted, P. et al. Modular, efficient and constant-memory single-cell RNA-seq preprocessing. Nat. Biotechnol. 39, 813–818 (2021).

71. Virshup, I., Rybakov, S., Theis, F. J., Angerer, P. & Wolf, F. A. anndata: Access and store annotated data matrices. J. Open Source Softw. 9, 4371 (2024).

72. Wolf, F. A., Angerer, P. & Theis, F. J. SCANPY: large-scale single-cell gene expression data analysis. Genome Biol. 19, 15 (2018).

73. Traag, V. A., Waltman, L. & van Eck, N. J. From Louvain to Leiden: guaranteeing well-connected communities. Sci. Rep. 9, 5233 (2019).

74. McInnes, L., Healy, J., Saul, N. & Großberger, L. UMAP: Uniform Manifold Approximation and Projection. J. Open Source Softw. 3, 861 (2018).

75. Squair, J. W. et al. Confronting false discoveries in single-cell differential expression. Nat. Commun. 12, 5692 (2021).

76. Crowell, H. L. et al. muscat detects subpopulation-specific state transitions from multi-sample multi-condition single-cell transcriptomics data. Nat. Commun. 11, 6077 (2020).

77. Robinson, M. D., McCarthy, D. J. & Smyth, G. K. edgeR: a Bioconductor package for differential expression analysis of digital gene expression data. Bioinformatics 26, 139–140 (2010).

78. Robinson, M. D. & Oshlack, A. A scaling normalization method for differential expression analysis of RNA-seq data. Genome Biol. 11, R25 (2010).

79. Haynes, W. Benjamini-Hochberg Method. in Encyclopedia of Systems Biology 78 (Springer New York, New York, NY, 2013). doi:10.1007/978-1-4419-9863-7_1215.

80. Heinz, S. et al. Simple combinations of lineage-determining transcription factors prime cis-regulatory elements required for macrophage and B cell identities. Mol. Cell 38, 576–589 (2010).

81. Mudge, J. M. et al. GENCODE 2025: reference gene annotation for human and mouse. Nucleic Acids Res. 53, D966–D975 (2025).

