## Supplementary Table legends and Supplemetary Figures for "Axolotl regeneration reveals a dormant cis-regulatory grammar conserved across vertebrate genomes"

### Supplementary Tables

**Supplementary Table 1 | Regeneration-prior and motif-grammar summaries**

Summary table of the axolotl regeneration prior and motif grammar used for cross-species DREC discovery. Rows report stage-level ATAC peak counts, dynamic and constitutive peak summaries, ATAC-RNA integration counts, motif-family enrichment summaries from day-7-versus-0-h expression-linked opening peaks, and *de novo* motif records from the day-7 expression-linked analysis. The table also records insertion eligibility and selection of the three *de novo* motifs used in the insertion analysis. The selected motifs are denovo_035 (HOMER header rank 24; CATGCAGATAAA; closest known match, FUS3), denovo_026 (header rank 15; TAGAATTGTGAA; closest known match, ZNF35), and denovo_033 (header rank 22; GCACTGAGAAAG; closest known match, PRDM1). Fields not applicable to a given section are left blank.

**Supplementary Table 2 | Candidate-construction quality-control summaries**

Quality-control summary table for ortholog mapping, motif-positive pre-candidate construction and AlphaGenome compatibility filtering. The table reports the transition from 181 axolotl regeneration-prior genes to 115 primary ortholog rows, 658 motif-positive pre-candidates and 649 AlphaGenome-scored DREC candidates, together with the nine-candidate exclusion audit, chromatin-class counts, score-bin poised/polycomb summaries and top-ranked candidate score audit. Fields not applicable to a given section are left blank.

**Supplementary Table 3 | Final dual-species DREC catalog**

Catalog of 649 human and mouse dormant regeneration enhancer candidates (DRECs), comprising 199 human (hg38) and 450 mouse (mm10) candidate intervals. For each candidate, the table reports locus identifiers, species, genome build, candidate-interval coordinates, linked genes, motif-count and motif-source fields, AlphaGenome-derived score fields including the DREC dormancy score and its bivalency/dormancy support components, predicted chromatin-class labels, screening priority, public-data support fields, conservation fields and validation-rank fields. The start, end and coord fields (candidate_interval_length_bp = end − start; median 220,625 bp, maximum 1,940,410 bp) denote motif-positive candidate intervals generated by chaining motif hits separated by no more than 128 bp within ortholog-centered search windows; they do not denote the full ortholog-centered search windows themselves. The midpoint-centered AlphaGenome input and scoring coordinates are provided separately as alphagenome_input_coord and alphagenome_scoring_coord, corresponding to the 131,072-bp input sequence and central 4,096-bp scoring interval used for AlphaGenome summarization. Candidates satisfying none of the five chromatin-class definitions are labeled unclassified. Multiple linked genes are separated by semicolons in the table. AlphaGenome scoring was performed over a fixed 4,096-bp window centered on midpoint (see alphagenome_scoring_start/alphagenome_scoring_end), independent of the candidate-interval length.

**Supplementary Table 4 | External support and sensitivity analyses**

Candidate-level and assay-level external support summaries for the DREC atlas. The table includes AlphaGenome-to-measured epigenome concordance, candidate-versus-background epigenomic support, strict TSS/GC-matched background sensitivity, GWAS Catalog overlap summaries, UCSC phyloP/phastCons conservation support, mouse postnatal chromatin-dynamics support and mouse injury/scATAC support. Public mouse developmental support includes GSE123867, and mouse injury/scATAC support includes GSE153479.

**Supplementary Table 5 | Locus-level summaries of motif-level *in silico* mutagenesis**

Locus-level summaries for two predefined ISM analysis sets: the human-mouse common 18-locus set and the extended top-dormant 20-locus set. For the human-mouse common set, 78,931 unique native motif targets were evaluated in triplicate, yielding 236,793 replicate-level rows. For the top-dormant 20-locus set, 89,720 unique native motif targets were evaluated in triplicate, yielding 269,160 replicate-level rows. n_disruption_rows reports replicate-level rows, n_native_motif_targets reports unique native motif targets, and replicate_aggregation_method records that target-level effects were summarized using the maximum across three independent random-replacement replicates. The three inserted motifs are denovo_035 (CATGCAGATAAA; closest known match, FUS3), denovo_026 (TAGAATTGTGAA; closest known match, ZNF35), and denovo_033 (GCACTGAGAAAG; closest known match, PRDM1). DREC dormancy scores in this table are rounded to three decimal places; full-precision values are provided in **Supplementary Table 3**.

### Supplementary Figure Legends

**Supplementary Figure 1 | Temporal organization of chromatin accessibility and prioritization of regeneration-associated peak-gene links.**

**(a)** Heatmap of dynamic ATAC-seq peaks across eight stages of axolotl limb regeneration: 0 h, 3 h, and days 1, 3, 7, 14, 22, and 33. Rows represent individual dynamic peaks (15,132 total) and columns represent sampling time points. Color scale indicates the row-wise *Z* score of chromatin-accessibility signal, revealing distinct temporal patterns of transient, delayed, and sustained accessibility, including a prominent accessibility increase at day 7. Peaks were classified as dynamic if absent at 0 h and detected in at least two post-injury stages.

**(b)** Principal component analysis (PCA) of stage-resolved ATAC-seq profiles. PC1 and PC2 explain 86.5% and 2.6% of the total variance, respectively. Points represent individual regeneration stages, and arrows indicate their temporal order across the regeneration time course. The separation of later regenerative stages (days 14, 22, and 33) from pre-injury and early-response states along PC1 reflects the progressive chromatin reorganization accompanying blastema maturation.

**(c)** Volcano plot of pseudobulk gene-expression changes at day 7 relative to the uninjured 0-h state. The horizontal axis indicates $\log_2$ fold change (7 d vs 0 h) and the vertical axis indicates $-\log_{10}$(adjusted *P* value). Genes linked to ATAC-seq peaks by the peak-gene integration analysis are highlighted in red, whereas other genes are shown in gray. Dashed lines indicate FDR $< 0.05$ and $|\log_2$ fold change$| = 1$ shown for reference; only the FDR threshold was applied, and ten selected highly significant linked genes are outlined in black.

**(d)** Top-ranked ATAC-RNA peak-gene associations according to their support scores. Each bar represents an individual gene-peak pair, labeled by the linked gene and peak identifier. Higher scores indicate peak-gene associations supported more strongly by the multi-omic integration workflow. The support score (range 0–6) integrates ATAC-seq and RNA-seq effect-size magnitude, accessibility-expression correlation, concordant directionality between modalities, peak annotation class, and RNA differential-expression significance.

**Supplementary Figure 2 | Temporal accessibility-expression relationships and motif composition of day-7 expression-linked regulatory elements.**

**(a)** Stage-resolved ATAC-seq and pseudobulk RNA-seq profiles for genes associated with top-ranked peak-gene pairs. Rows represent the top 23 ATAC-RNA linked genes ranked by support score, and columns show ATAC-seq and RNA-seq signals across the eight regeneration stages. Color scale indicates the row-wise $Z$ score within each data modality. The heatmap illustrates gene-specific temporal relationships between chromatin accessibility and transcription, including both concordant and temporally offset changes. Genes are ordered by decreasing support score, using the same ranking criterion as in **Supplementary Fig. 1**d (which displays the top 20 individual gene-peak pairs and therefore comprises a different number of entries).

**(b)** Known transcription factor motifs enriched in day-7 accessible peaks linked to gene-expression changes, identified by HOMER known-motif analysis against a background of all master ATAC-seq peaks. Bars indicate motif-enrichment significance, expressed as $-\log_{10} P$, and bar color denotes the percentage of target peaks containing the corresponding motif. The motif names in the figure are derived from HOMER. Enriched motifs include AP-2γ, HOXA13, HIC1, HOXC13, and HOXA9, representing a developmental- and signal-responsive transcription factor repertoire consistent with the broader regenerative grammar defined in **Fig. 2g**.

**(c)** Sequence logos of the five most significantly enriched known motifs in day-7 expression-linked accessible peaks: AP-2γ, HOXA13, HIC1, HOXC13, and HOXA9. Enrichment scores ($-\log_{10} P$) are shown beside each motif name. Logos were generated from HOMER motif files and represent the position weight matrix of each factor's binding preference. The motif names in the figure are derived from HOMER.

**(d)** Sequence logos of five representative *de novo* motifs from the HOMER *de novo* overview of day-7 expression-linked accessible peaks. The score above each logo is the HOMER overview enrichment score and is on a different scale from the source_log_p_support statistic used to rank insertion eligibility (**Supplementary Table 1**); the two should not be compared directly. Motif 1 is the reverse-complement auxiliary record of motif 2, motif 3 is the reverse-complement auxiliary record of motif 4, and motif 5 is the reverse-complement auxiliary record of a primary motif not shown in the panel. Thus, the five displayed records represent three underlying primary motif classes. The insertion analysis used a separate prespecified eligible primary-motif subset; the selected motifs were denovo_035, denovo_026 and denovo_033 (HOMER header ranks 24, 15 and 22), none of which is displayed in this panel. Selection was performed without reference to AlphaGenome insertion outcomes.

**Supplementary Figure 3 | Mouse external validation.**

**(a)** Candidate-level mouse external H3K27me3 validation stratified by AlphaGenome-predicted chromatin class. Each point represents one mDREC candidate and reports $\log_2$ candidate-versus-matched-control H3K27me3 signal summarized across public mouse H3K27me3 datasets. Chromatin classes are non-exclusive, so a candidate with multiple predicted labels can appear in more than one class-stratified group. Asterisks indicate $P < 1 \times 10^{-4}$ for one-sided candidate-greater comparisons against matched controls.

**(b)** Mouse poised/polycomb contrast. Each mDREC is counted once, with candidates carrying either a poised-enhancer-like or polycomb-marked label assigned to the poised/polycomb group and all remaining candidates assigned to the other-DREC group. The panel tests whether external H3K27me3 support is concentrated in the predicted poised/polycomb subset. Asterisks indicate $P < 1 \times 10^{-4}$ for the one-sided poised/polycomb-greater comparison.

**(c)** Mouse postnatal H3K27ac dynamics at prioritized poised/polycomb mDREC loci. Bars show the change in H3K27ac signal from postnatal day 1 to postnatal day 8 (P8–P1) for prioritized poised/polycomb mDREC loci, ranked by H3K27ac change. Negative values indicate postnatal loss of active-enhancer signal, and the dashed line indicates the median change across the plotted loci.

**Supplementary Figure 4 | Human external validation.**

**(a)** Candidate-level human external H3K27me3 validation stratified by AlphaGenome-predicted chromatin class. Each point represents one hDREC candidate and reports $\log_2$ candidate-versus-matched-control H3K27me3 signal summarized across public human H3K27me3 datasets. Chromatin classes are non-exclusive, so a candidate with multiple predicted labels can appear in more than one class-stratified group. Asterisks indicate $P < 1 \times 10^{-4}$ for one-sided candidate-greater comparisons against matched controls.

**(b)** Human poised/polycomb contrast. Each hDREC is counted once, with candidates carrying either a poised-enhancer-like or polycomb-marked label assigned to the poised/polycomb group and all remaining candidates assigned to the other-DREC group. The panel tests whether external H3K27me3 support is concentrated in the predicted poised/polycomb subset. Asterisks indicate $P < 1 \times 10^{-4}$ for the one-sided poised/polycomb-greater comparison.

**(c)** Human GWAS annotation-overlap analysis. hDREC candidates and matched background controls were overlapped with GWAS Catalog associations grouped into a composite cardiac/muscle/neurologic set and separate cardiac, muscle and neurologic sets using predefined text filters. Bars show the percentage of candidate and background intervals overlapping annotations within the analyzed window. The composite set remained significant after Bonferroni correction across the four trait sets (adjusted $P = 9.19 \times 10^{-3}$), whereas the cardiac, muscle and neurologic sets did not.

**Supplementary Figure 5 | Locus selection and baseline chromatin state of the *in silico* motif perturbation across human-mouse matched DREC loci.**

**(a)** DREC dormancy score for the nine human-mouse ortholog gene neighborhoods analyzed by *in silico* motif perturbation, comprising 18 matched loci in total. Each pair of bars shows the human (blue) and mouse (orange) locus scores. Loci were selected for perturbation on the basis of requiring a high DREC dormancy score in both species and the availability of a confident one-to-one ortholog assignment. Human and mouse scores are shown side by side for descriptive visualization of the selected ortholog-associated pairs; their raw magnitudes are not interpreted as quantitatively comparable between species because PRO-cap contributed only to the human score.

**(b, c)** Baseline (wild-type) predicted epigenomic profiles of the perturbed loci for human (b) and mouse (c). Rows represent individual loci ranked by DREC dormancy score, corresponding to the nine orthologous gene neighborhoods selected for perturbation analysis. Columns represent seven predicted tracks: H3K4me1, H3K27me3, H3K27ac, ATAC, DNase, CAGE, and PRO-cap. Color scale indicates the *Z* scored predicted signal intensity. Human profiles include seven tracks. Mouse profiles include the six available tracks; PRO-cap predictions were unavailable and are shown in gray as not scored. Co-enrichment of H3K4me1 and H3K27me3 together with relative suppression of H3K27ac, chromatin-accessibility, and transcriptional-output tracks recapitulates the bivalent dormancy signature described in **Fig. 3f**. Per-locus numeric values are given in **Supplementary Tables 3 and 5**.

**Supplementary Figure 6 | Predicted effects of *in silico* motif perturbation at**

**human-mouse matched DREC loci.**

**(a)** Relationship between predicted motif-disruption importance and predicted opening-gain score across the top 25 non-redundant disruptions, all of which occur at human loci. Each point represents one motif instance; the highest-importance disruption at each locus is summarized in **Supplementary Table 5**. Stronger disruptions tend to combine H3K27me3 loss with an increase in predicted chromatin opening, consistent with a coordinated transition from the dormant bivalent state toward an enhancer-like chromatin configuration.

**(b)** Predicted change in histone-modification signal (Δ = variant minus wild-type) at the 15 strongest motif disruptions ranked by disruption importance. For each disruption, filled circles show the predicted change in H3K27me3 (poised, navy) and H3K27ac (active, pink), connected by a line. The locus and disrupted motif are indicated on the vertical axis. Repeated locus–motif labels denote distinct motif instances at different target coordinates. Disruptions consistently decrease H3K27me3 and increase H3K27ac, illustrating a predicted shift from a dormant poised/repressed chromatin state toward an active enhancer-like state.

**(c)** Predicted wild-type and variant signal tracks (H3K27me3, H3K27ac, ATAC, and DNase) for a representative native motif disruption (human *PIK3R2*, EAR2). Tracks are shown across a ±10-kb window centered on the candidate locus midpoint. In each track, the wild-type prediction is shown as a gray dashed line and the variant prediction as a solid colored line, with the signal difference between variant and wild-type reflecting the predicted chromatin remodeling at each position. The vertical orange line marks the disrupted motif position. In this panel, disruption of the EAR2 motif at the human *PIK3R2* locus produces a pronounced predicted decrease in H3K27me3 accompanied by gains in H3K27ac, ATAC, and DNase signal, consistent with dormancy destabilization.

**(d)** Maximum motif-disruption importance at each locus. Each bar represents the largest disruption importance score among all native motif instances tested at that locus, for human (blue) and mouse (orange) candidates. Disruption importance is defined as the predicted shift away from the dormant bivalent state upon replacement of the motif sequence with a randomized sequence of equal length. For each native motif target, the maximum value across three independent random-replacement replicates was retained, and each plotted bar shows the maximum target-level value at that locus.

**(e)** Maximum *de novo* motif insertion opening gain at each locus, for human (blue) and mouse (orange) candidates. Opening gain reflects the predicted shift toward an accessible, enhancer-like chromatin state following insertion of a FUS3-like, PRDM1-like, or ZNF35-like *de novo* motif at the locus center; the value shown is the maximum across the three motif types. Most loci show little predicted opening gain, with the strongest response at the mouse *Elf3–Lad1* locus. Several loci (human *CDKN1B*, *CKB*, *TRIM56* and *USP2*, and mouse *Cinp–Ckb*) show slightly negative values, indicating a predicted reduction rather than gain of chromatin opening, consistent with context-dependent suppression of accessibility upon motif insertion at these sites.

**(f)** Predicted wild-type and variant signal tracks (H3K27me3, H3K27ac, ATAC, and DNase) for a representative *de novo* motif insertion (mouse *Elf3–Lad1*, FUS3-like). Tracks are shown across a ±10-kb window centered on the candidate locus midpoint.

In each track, the wild-type prediction is shown as a gray dashed line and the variant prediction as a solid colored line, with the signal difference between variant and wild-type reflecting the predicted chromatin remodeling at each position. The vertical orange line marks the motif insertion site. In this panel, *de novo* FUS3-like motif insertion at the mouse *Elf3–Lad1* locus drives a predicted gain in H3K27ac and chromatin accessibility with modest H3K27me3 reduction, consistent with partial chromatin opening.

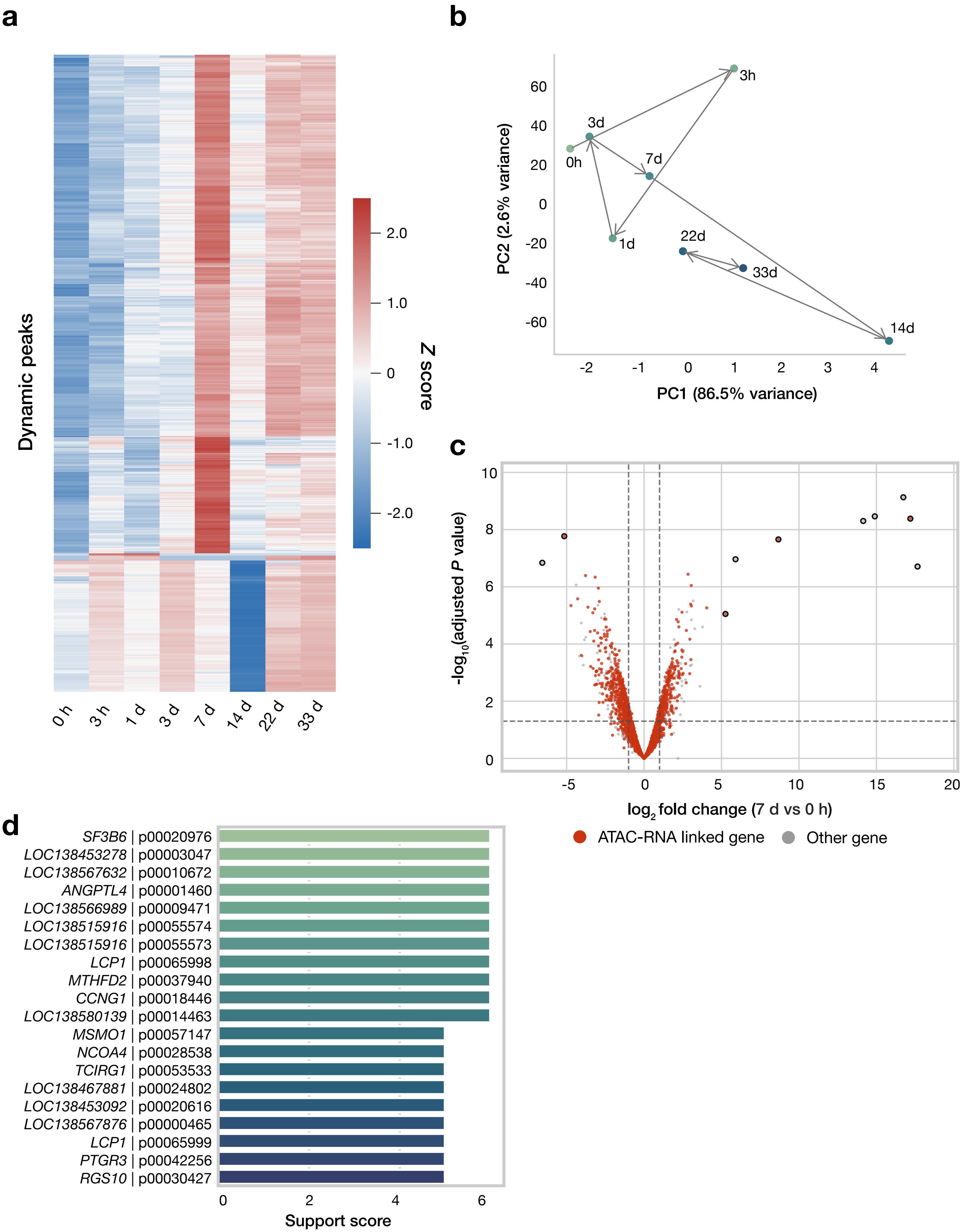


**Fujiwara *et al.* Supplementary Figure 1**

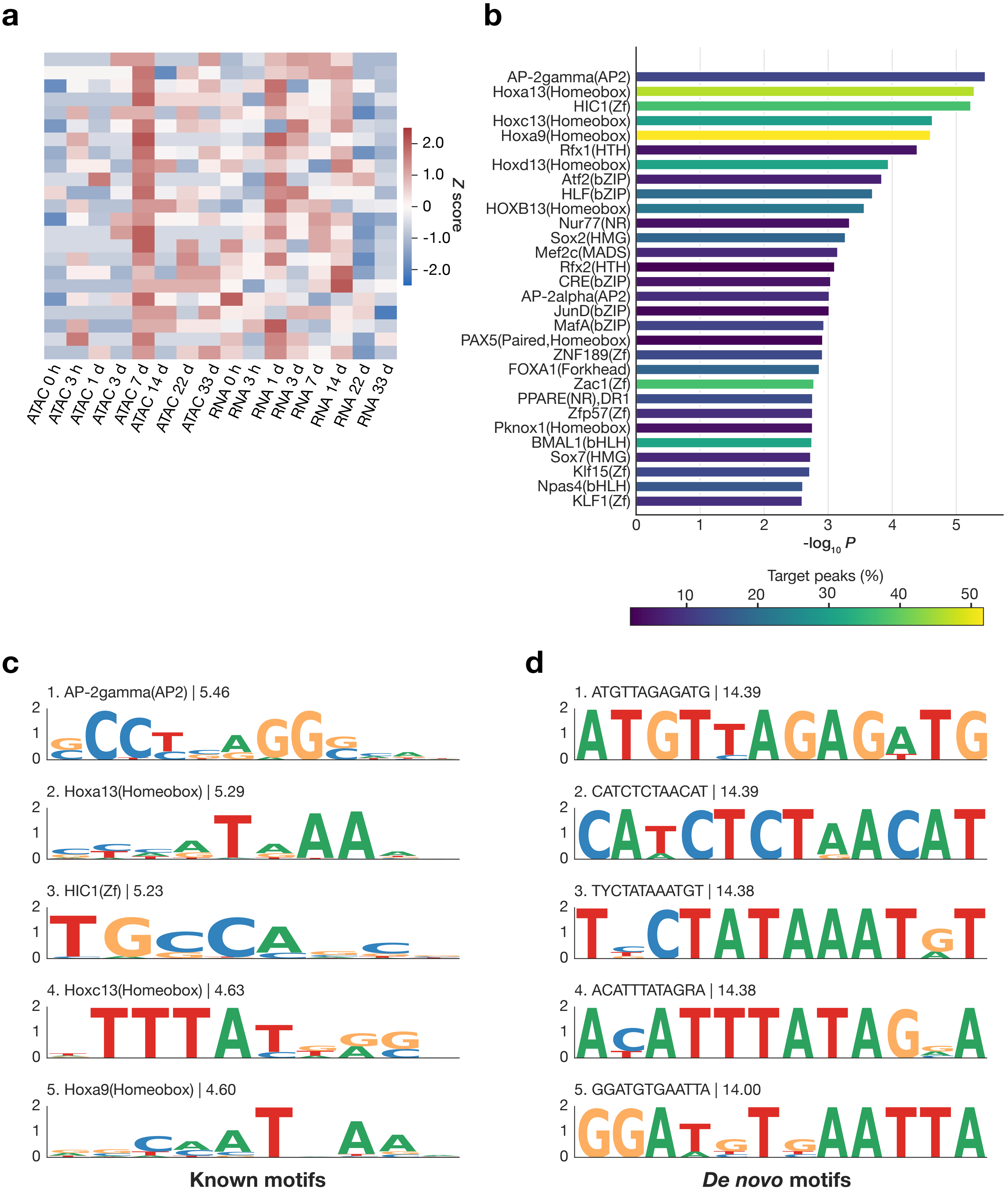


**Fujiwara *et al.* Supplementary Figure 2**

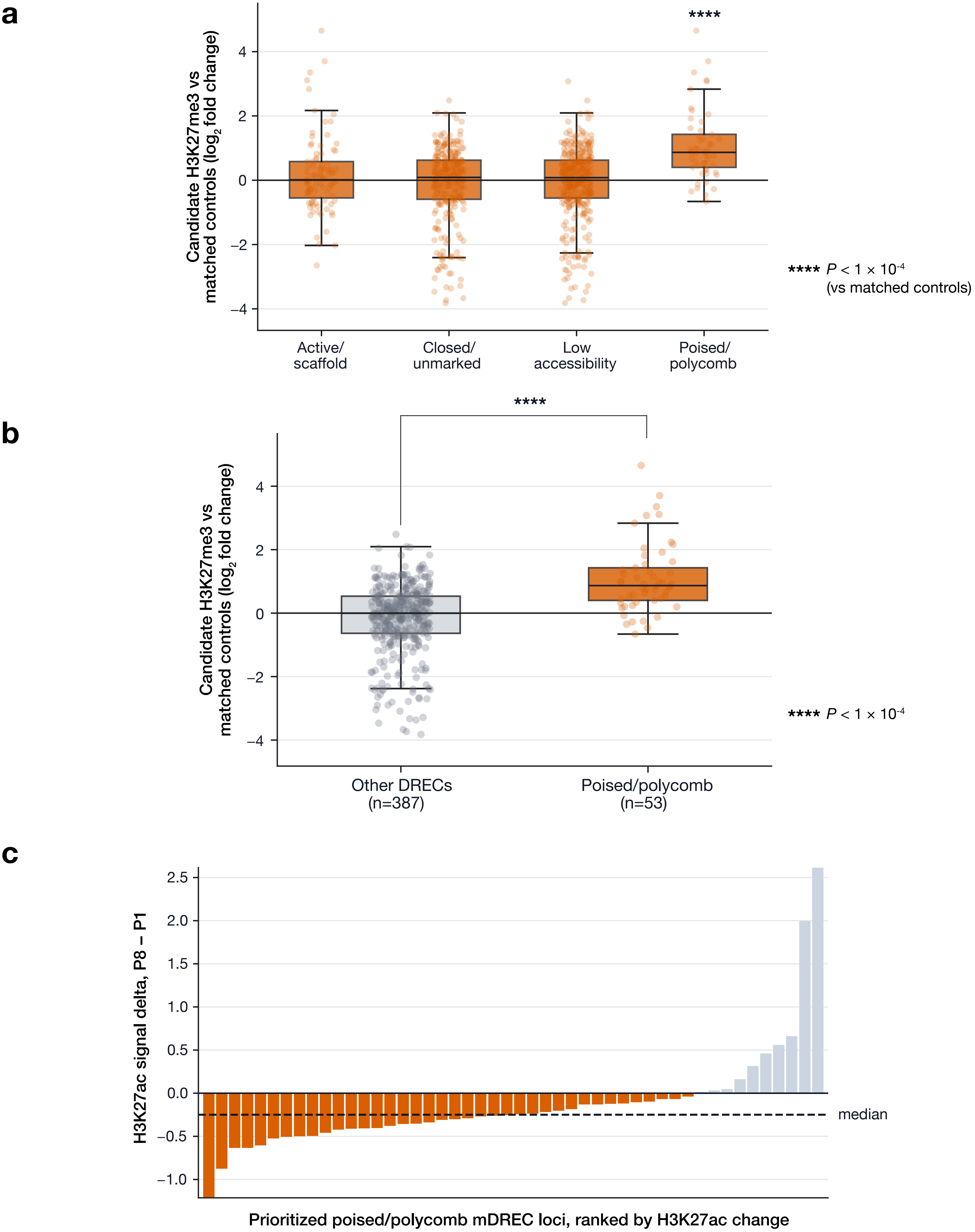


**Fujiwara *et al.* Supplementary Figure 3**

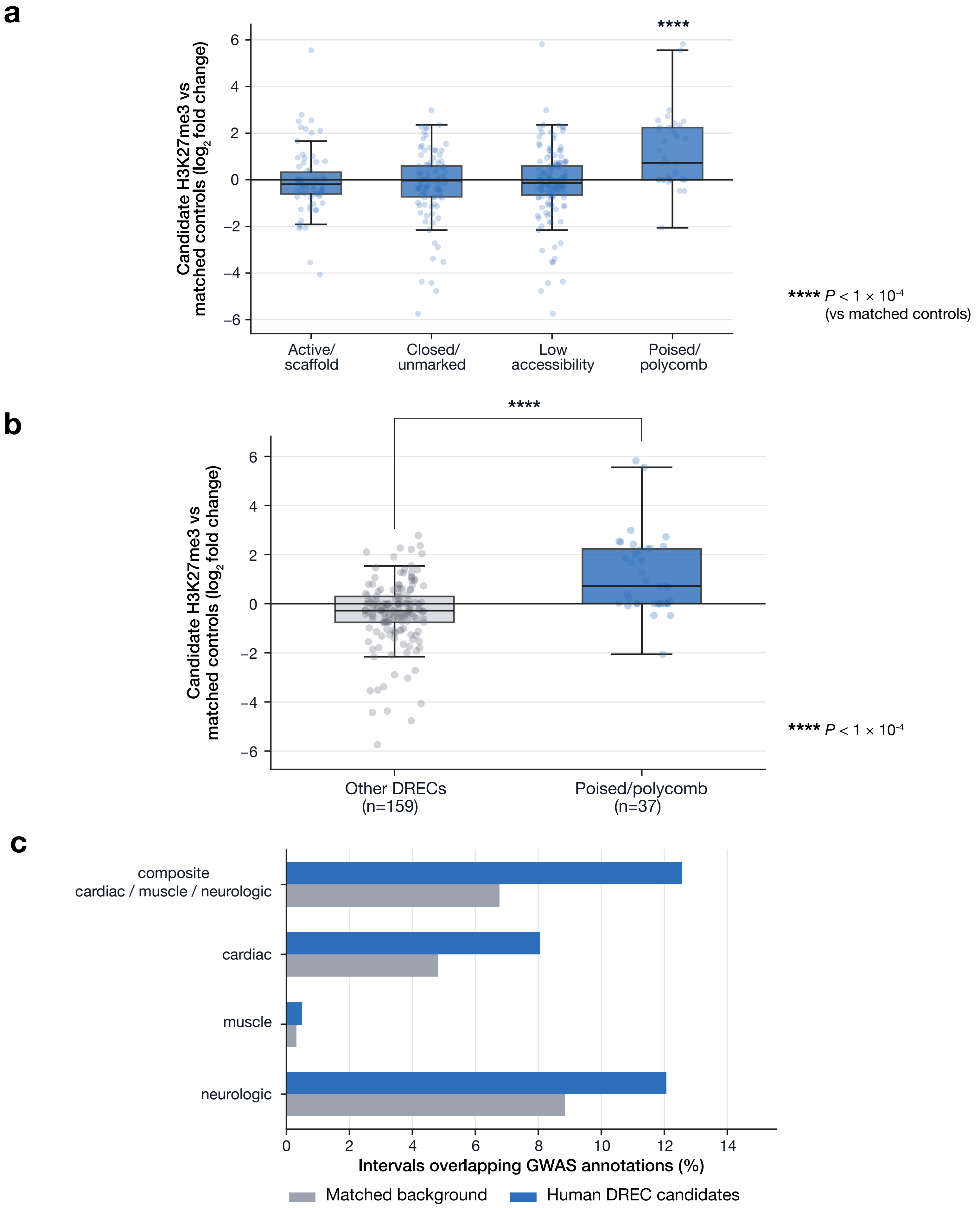


**Fujiwara *et al.* Supplementary Figure 4**

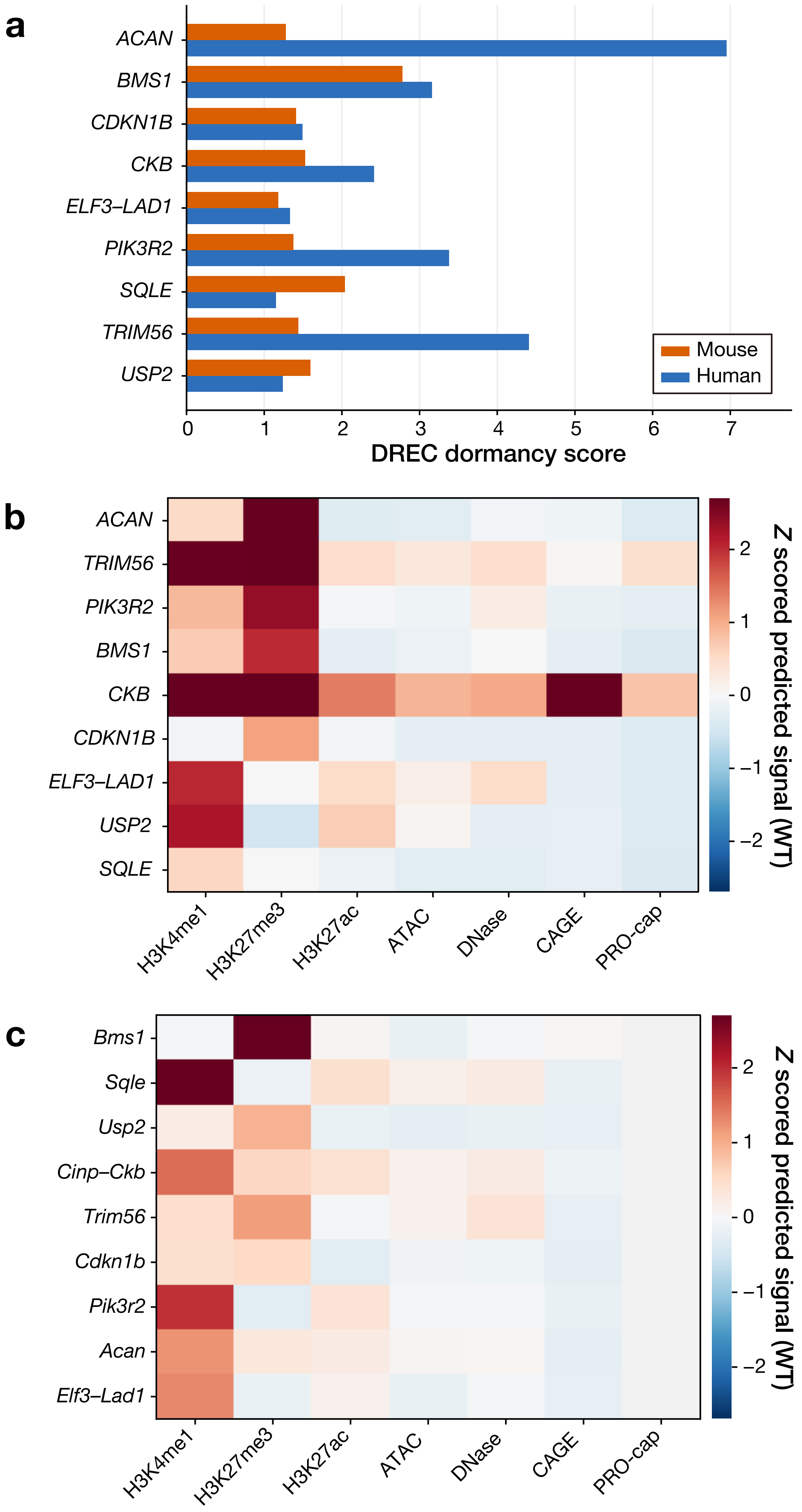


**Fujiwara *et al.* Supplementary Figure 5**

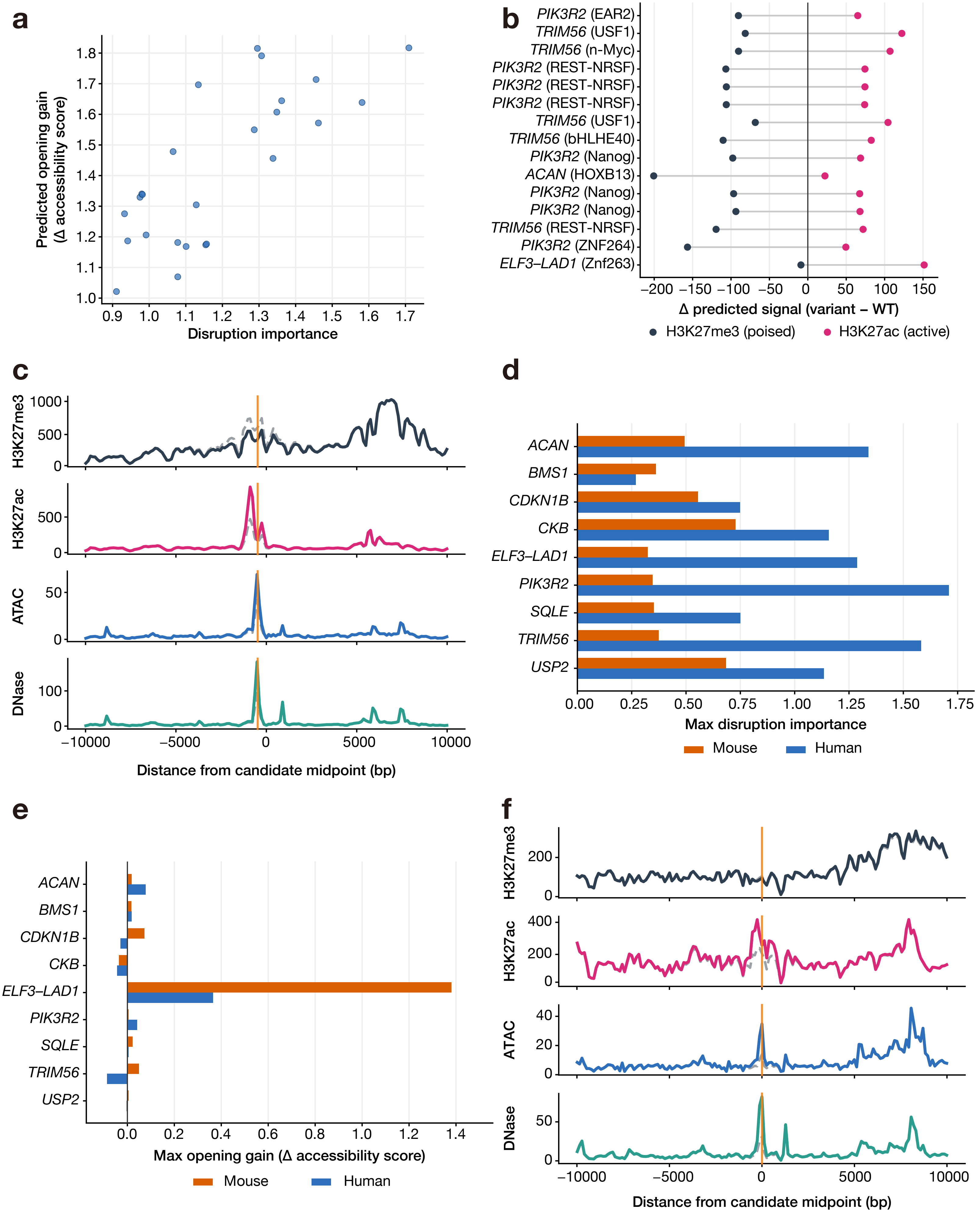


**Fujiwara *et al.* Supplementary Figure 6**
